# Rapid High-Throughput Analysis of Bovine Skeletal Muscle Fiber Morphology via Automated Fluorescent Microscopy and MuscleBos software

**DOI:** 10.64898/2025.12.18.695234

**Authors:** Hamood Rehman, Kyrstin M. Gouveia, Rebecca K. Coombe, Jacquelyn P. Boerman, J. Alex Pasternak, James F. Markworth

## Abstract

Skeletal muscle tissue is comprised of many individual muscle cells (myofibers) that can be classified as different types based on their morphology, histochemistry, enzymatic reactivity, and biochemical characteristics. One of the most common methods of classification of muscle fiber type relies on the local expression of specific myosin heavy chain (MyHC) isoforms. Adult mammalian muscle fibers are generally categorized into four major types including I, IIA, IIX, and IIB. However, the distribution of these muscle fiber types varies across both different species and muscle groups within species, influencing muscle function and physiological responses. In bovine species, skeletal muscle plays a critical role in determining *in-vivo* metabolic physiological processes and impacting post-harvest meat quality traits. Immunostaining methods using isoform-specific MyHC antibodies have been widely adopted to characterize muscle fiber morphology. However, manual capture and analysis of immunofluorescent images of muscle fiber type staining is time consuming, labor-intensive, and potentially susceptible to investigator bias. To address these limitations, we established and validated a high-throughput method for the analysis of bovine muscle fiber morphology that combines automated fluorescent microscopy with high-content image analysis using a customized version of the MuscleJ plugin for FIJI/ImageJ that we named MuscleBos. This refined method enables rapid quantitative characterization of muscle fiber type profile and fiber type-specific myofiber cross-sectional area in bovine skeletal muscle tissue cross-sections. This methodology should enable valuable deeper insights into future studies of muscle composition in bovine species and its impact on *in vivo* animal physiology and meat science.

## Background

Skeletal muscle is heterogeneous tissue composed of many individual multinucleated contractile muscle cells termed myofibers that can be classified into various types based on their morphology, histochemistry, enzymatic reactivity, and biochemical characteristics (1). One of the most common methods of classification of muscle fiber type involves identifying the expression of specific myosin heavy chain (MyHC) isoforms (2). In adult mammals, skeletal myofibers are typically categorized into four main types including type I, IIA, IIX, and IIB (3). The presence of type IIB fibers generally appears restricted to small mammals (4). Type I fibers are relatively red in color, slow-twitch, fatigue resistant, and have a high oxidative capacity (3). Type IIA fibers are relatively white in color and fast-twitch, while still being relatively fatigue-resistant and oxidative. Type IIB fibers when present are also white and fast-twitch but are distinguished from type IIA fibers by their relatively greater fatigability and higher glycolytic activity (5). Finally, type IIX fibers are generally considered intermediate between type IIA and type IIB fibers in their properties (6). In addition to these pure fiber types, hybrid fibers are frequently observed in healthy adult muscle tissue (7).

Muscle fiber type composition in cattle is an important determinant of meat tenderness and flavor (4, 8–10). The fiber type composition of bovine muscle differs most notably between muscle groups with distinct physiological demands (11, 12). For example, the deep *psoas major* muscle (tenderloin) has a relatively higher proportion of type I muscle fibers when compared to the *longissimus dorsi* (e.g., loin) and *semitendinosus* (e.g., eye of round) in which type II fibers predominate (13–16). Muscle fiber type profile can also switch in response to altered functional demands (17). Fiber type composition in beef cattle can be influenced by age (18), breed (19), genetic selection (20, 21), method of housing (22), biological sex (23), nutritional state (24–26), and administration of anabolic steroids and/or beta adrenergic agonists (27–32). While fewer studies are available in dairy cows, dietary supplementation with rumen protected niacin from 21 d before expected calving until 3-weeks postpartum resulted in a fiber type shift from type IIX towards type IIA in the *semitendinosus* muscle (33).

Techniques have been developed to identify muscle fibers of various types, each with its advantages and limitations (34). Among these, immunohistochemical staining with fiber type specific primary antibodies against MyHC isoforms has become the most widely used method (5). In the last decade there has been a surge in the availability of purpose-built semi or fully automated software options for the image analysis of immunofluorescent muscle cross-sections (35–47). With a few exceptions (39, 46), the majority of these software options can also characterize muscle fiber type (35–38, 40–45, 47). However, all these software were originally developed and validated using tissue samples obtained from mice (35–47), rats (41, 42, 44), or humans (35, 48). In contrast, image analysis is generally still performed manually in studies of bovine muscle histology (e.g., 15, 28, 29, 33, 49, 50–52). Such manual image analysis is labor-intensive, time-consuming, and potentially prone to investigator bias. To address these limitations some commercially available software such as Noesis Visilog (53) and Agilent Gen5 (54–56) have been applied to the analysis of immunofluorescent staining of bovine muscle samples. The only freely available software that has been specifically validated for use on bovine muscle samples is the deep learning-based tool MyoV (57). However, MyoV is designed for the analysis of brightfield images of hematoxylin and eosin (H&E)-staining and as such cannot identify muscle fiber type profile or quantify fiber type-specific myofiber size (57).

In the current study we established and validated a rapid high-throughput method for analysis of bovine myofiber morphology, muscle fiber type, and fiber type-specific myofiber size. This method combines automated immunofluorescent microscopy and high-content image analysis using MuscleBos software, a new customized bovine-specific version of the open source and freely available MuscleJ plugin in ImageJ/FIJI (45). Ultimately, this novel methodology should enable valuable deeper insights into future studies of skeletal myofiber morphology in bovine species and its impact on *in vivo* animal physiology and meat science.

## Materials and Methods

### Animals and housing

Data for this study were collected, and procedures were performed at the Purdue University Animal Sciences Research and Education Center (ASREC) dairy unit from September 2022 through February 2023 as part of a larger study (58). Muscle biopsy samples were collected from n=48 multiparous Holstein dairy cows that had completed 2.8 ± 1.0 (mean ± SD) lactations, with previous 305-day milk yield of 11,382 ± 1,338.3 kg, and body weight of 749 ± 72.2 kg at the time of enrollment which was 42 days before expected calving. For more information regarding the animals and housing please refer to Gouveia et al. (2024) (58).

### Skeletal Muscle Biopsies

At approximately 21 d before expected calving and 21 days in milk, a biopsy sample was obtained from the *longissimus dorsi* muscle of each cow by the same trained individual. Cattle were restrained in a livestock handling chute, hair was clipped from the 9th to 13th rib, and the underlying skin cleaned thoroughly. The area was cleaned with 70% isopropyl alcohol, scrubbed with povidone-iodine, and then cleaned once more with 70% isopropyl alcohol. A local anesthetic (2% lidocaine, 5 mL) was then injected into the intercostal area. Under aseptic conditions, an incision was made with a #10 scalpel blade through hide and adipose tissue to expose the underlying muscle tissue. A 5 mm Bergstrom needle was then used to collect muscle biopsy samples, cutting in a circular pattern to ensure a representative sample of the area. The collected muscle tissue was washed in a beaker containing sterile 1 × phosphate buffered saline (PBS). The tissue sample obtained was confirmed as muscle via a float test, in which muscle tissue sinks while adipose tissue floats. Following sample collection, the surgical area was once again cleaned with 70% isopropyl alcohol gauze before being patted dry with sterile dry gauze. After drying, the incision was closed using tissue adhesive and sprayed with an aerosol bandage. The incision site and animal were monitored for a week following the procedure with no complications or health issues occurring from the operation.

### Immunofluorescence analysis of bovine muscle fiber type

A portion of each bovine *longissimus dorsi* muscle biopsy sample was oriented longitudinally on a plastic support, covered with optimal cutting temperature (OCT) compound (Fisher Scientific, 23-730-571), and rapidly frozen in isopentane cooled on liquid nitrogen. OCT embedded muscle biopsy samples were stored at -80°C until further analysis. Muscle tissue cross-sections (10 µm) were cut in a cryostat (Leica CM1950) at -20°C and adhered to SuperFrost Plus slides (Fisher Scientific, 12-550-15). Slides were air dried at room temperature, tissue sections were encircled with hydrophobic PAP pen (Vector, H-4000) and then blocked for 1 h at room temperature in 10% goat serum (GS) (Invitrogen, 10000C) prepared in PBS. Slides were incubated overnight in a humidified chamber at 4°C with primary antibodies diluted in blocking buffer. Primary antibodies tested included those against dystrophin [Developmental Studies Hybridoma Bank (DSHB), MANDYS1(3B7)s, MIgG2a, 1:10], Laminin 1 + 2 (Abcam, ab7463, Rabbit, 1:100), MyHC type I (DSHB, BA-D5c, MIgG2b, 1:100), MyHC type IIA (DSHB, SC-71c, MIgG1, 1:100), MyHC type IIX (DSHB, 6H1s, MIgM, 1:10), and all but type MyHC IIX (DSHB, BF-35c, MIgG1, 1:100). The following day slides were washed three times for five minutes each in PBS prior to incubating with appropriate Alexa Fluor conjugated secondary antibodies (diluted 1:500 in PBS) including Goat Anti-Rabbit IgG (H+L) Alexa Fluor 350 (Invitrogen, A-11046), Goat Anti-Mouse IgG2a Alexa Fluor 647 (Invitrogen, A-21241), Goat Anti Mouse IgG1 Alexa Fluor 488 (Invitrogen, A-21121), Goat Anti-Mouse IgG2b Alexa Fluor 555 (Invitrogen, A-21147), and Goat Anti-Mouse IgM Alexa Fluor 647 (A-21238). Following further washing in PBS slides were mounted with coverslips using MOWIOL Fluorescence Mounting Medium and air dried at room temperature for 24 h.

### Immunofluorescent microscopy

Stitched mosaic images of the entire cross-section of each muscle biopsy sample were captured with a 10 × Plan Fluorite objective using an automated fluorescent microscope equipped with a motorized stage and DAPI, FITC, Texas Red, and CY5 filters operating in upright configuration (Echo Revolution). Pseudo-colors were applied to aid visual differentiation or various muscle fiber types. Stitched images of each individual fluorescent channel were exported as 16-bit TIFF file format and then merged as Multi-Channel TIFF images using ImageJ software prior to image analysis.

### High throughout image analysis with MuscleBos software

Multi-channel 16-bit TIFF images were analyzed using the MuscleJ 1.0.2 plugin for FIJI/ImageJ with custom modifications. MuscleJ is an open-source plugin for ImageJ/FIJI originally designed and validated for analysis of mouse skeletal muscle tissue sections (45). The MuscleJ plugin’s “Fiber morphology” function uses fluorescent images of myofiber border staining (e.g., dystrophin, laminin, or wheat germ agglutinin) to automatically segment, count, and measure the cross-sectional area (CSA) of each individual myofiber within a muscle cross-section. Additionally, MuscleJ includes a “Fiber type” function automatically quantifies the signal intensity of MyHC-specific antibody staining within each myofiber boundary relative to a threshold value to determine muscle fiber type profile and fiber type specific CSA. While initially designed and validated for use on mouse skeletal muscle tissue cross-sections, here we customized and validated MuscleJ 1.0.2 to enable rapid high-throughout the analysis of bovine muscle cross-sections. We named this new bovine-specific MuscleJ version MuscleBos.

### Statistical Analysis

Statistical analysis was performed in GraphPad Prism 10. The results obtained from MuscleBos both with and without type IIX fiber specific CSA exclusion thresholds applied were compared to those obtained via manual measurements of percentage muscle fiber type and fiber type specific myofiber CSA on the same source images via one-way analysis of variance (ANOVA) followed by pairwise Holm-Šídák post hoc tests. For application of the customized bovine MuscleJ plugin to the characterization of shifts in muscle fiber morphology between dairy cows with differing amounts of *longissimus dorsi* muscle mass throughout the transition period data was analyzed by two-way ANOVA followed by pairwise Holm-Šídák post hoc tests. Differences between groups were considered statistically significant at a cutoff of p<0.05.

## Results

### The bovine *longissimus dorsi* muscle lacks type IIB fibers

Muscle biopsy cross-sections were first stained with a primary antibody against laminin together with a cocktail of mouse monoclonal antibodies against MyHC type I (BA-D5), type IIA (SC-71), and type IIB (BF-F3) (**Fig. 1**). This primary antibody cocktail is commonly used in studies of rodent muscle fiber type (59). This staining revealed an apparent lack of type IIB fibers within the bovine *longissimus dorsi* muscle (**Fig. 1**). In rodent skeletal muscle, the SC-71 primary antibody reacts specifically with type IIA myofibers and does not stain IIX or IIB fibers (60–63). In contrast, we found that the SC-71 antibody appeared to react strongly with all non-type I (BA-D5^neg^) muscle fibers in bovine muscle biopsy cross-sections (**Fig. 1**).

**Figure 1:**
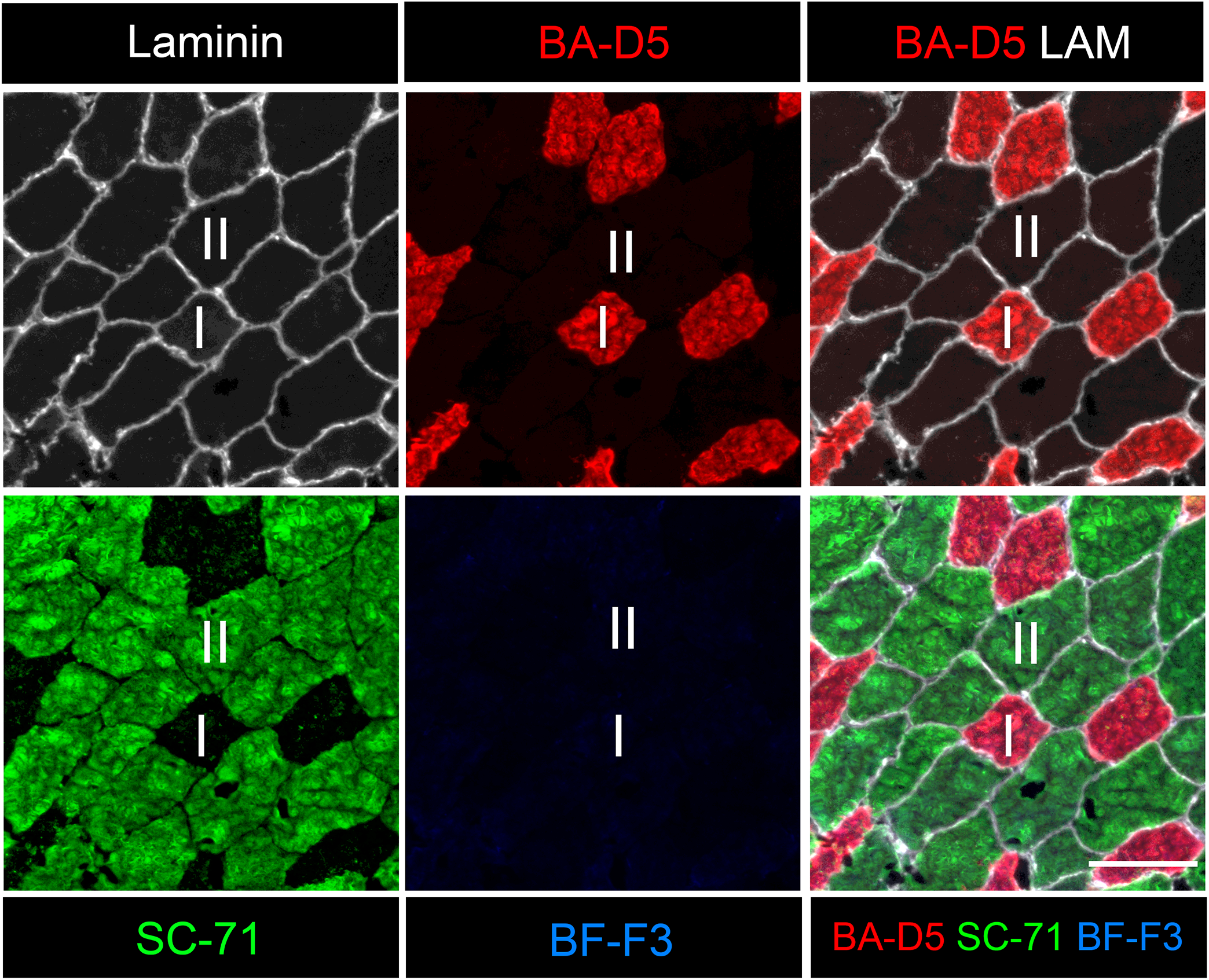
Identification of type I vs. II myofibers in bovine skeletal muscle: Cross-sections (10 µm) of bovine *longissimus dorsi* muscle biopsy samples were stained with a primary antibody against laminin (ab7463, Rabbit, 1:100) to label the myofiber boundaries, together with a cocktail of mouse monoclonal antibodies against MyHC type I (BA-D5c, MIgG2b. 1:100), MyHC type IIA (SC-71c, MIgG1, 1:100), and MyHC type IIB (BF-F3c, MIgM, 1:100). Primary antibody staining was visualized using Alexa Fluor-conjugated secondary antibodies (1:500) including Goat Anti-Rabbit Alexa Fluor 350 (to detect laminin), Goat Anti-Mouse IgG1 Alexa Fluor 488 (to detect SC-71), Goat Anti-Mouse IgG2b Alexa Fluor 555 (to detect BA-D5), and Goat Anti-Mouse IgM Alexa Fluor 647 (to detect BF-F3). Laminin, BA-D5, SC-71, and BF-F3 staining were pseudo colored white, red, green, and blue respectively. Scale bars 100 µm.

### Identification of type IIA and IIX myofibers in bovine skeletal muscle

To confirm the presence or absence of type IIX muscle fibers in these bovine *longissimus dorsi* samples, we stained serial bovine muscle cross-sections with a primary antibody against dystrophin in combination with a cocktail of mouse monoclonal antibodies against MyHC IIX (6H1) and all myosin isoforms except for type IIX (BF-35) (**Fig. 2**). This staining revealed many IIX myofibers as present based on both negative immunoreactivity with BF-35 and positive immunoreactivity for 6H1 (**Fig. 2**). Furthermore, type IIA and type IIX fibers stained equally brightly with the SC-71 antibody suggesting that these bovine fiber types cannot be reliably distinguished based on relative SC-71 staining intensity (**Fig. 2**).

**Figure 2:**
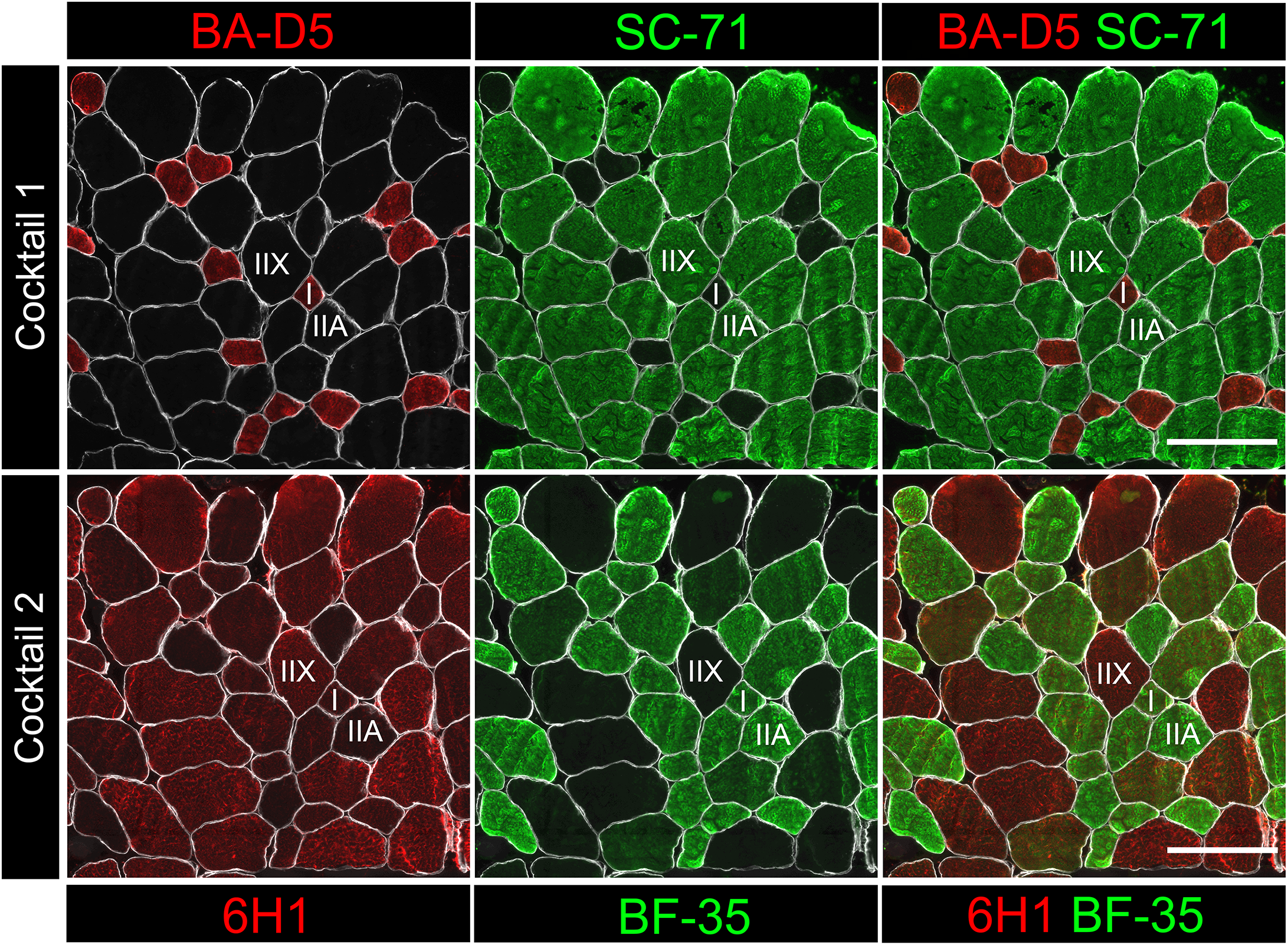
Distinguishing type IIA and IIX myofibers in bovine skeletal muscle: **A:** Cross-sections (10 µm) of bovine *longissimus dorsi* muscle biopsy samples were stained with a primary antibody against dystrophin [MANDYS1(3B7)s, MIgG2a, 1:10] to label the label the myofiber boundaries, together with a cocktail of mouse monoclonal primary antibodies against MyHC type I (BA-D5c, MIgG2b, 1:100) and MyHC type IIA (SC-71c, MIgG1, 1:100). Primary antibody staining was visualized using Alexa Fluor-conjugated secondary antibodies (1: 500) including Goat Anti-Mouse IgG2a Alexa Fluor 647 (to detect dystrophin), Goat Anti-Mouse IgG1 Alexa Fluor 488 (to detect SC-71), and Goat Anti-Mouse IgG2b Alexa Fluor 555 (to detect BA-D5). Dystrophin, BA-D5, and SC-71 staining was pseudo colored white, red, and green, respectively. **B:** Serial cross-sections from the same bovine *longissimus dorsi* muscle biopsy sample shown in panel A were stained with a primary antibody against dystrophin [MANDYS1(3B7)s, MIgG2a, 1:10] to label the label the myofiber boundaries in combination with a cocktail of primary antibodies against MyHC type IIX (6H1s, MIgM, 1:10) and all MyHC isoforms except for type IIX (BF-35c, MIgG1, 1:100). Primary antibody staining was visualized using Alexa Fluor-conjugated secondary antibodies (1:500) including Goat Anti-Mouse IgG2a Alexa Fluor 647 (to detect dystrophin), Goat Anti-Mouse IgG1 Alexa Fluor 488 (to detect BF-35), and Goat Anti-Mouse IgM Alexa Fluor 555 (to detect 6H1). Dystrophin, BF-35, and 6H1 staining were pseudo colored white, red, and green, respectively. Scale bars are 200 µm.

### Simultaneous identification of type I, IIA, and IIX fibers in bovine muscle

To establish a more reliable method of bovine muscle fiber type identification, we next tested different MyHC specific primary antibody combinations. We used a primary antibody against laminin together with a combination with monoclonal primary antibodies against MyHC type I (BA-D5), type IIX (6H1), and all but type IIX (BF-35) (**Fig. 3**). This primary antibody cocktail was able to identify type I myofibers (as BA-D5^pos^BF-35^pos^ cells), type IIA myofibers (as BA-D5^neg^BF-35^pos^ cells), and type IIX myofibers (as BF-35^neg^ 6H1^pos^ cells) on a single tissue section (**Fig. 3**).

**Figure 3:**
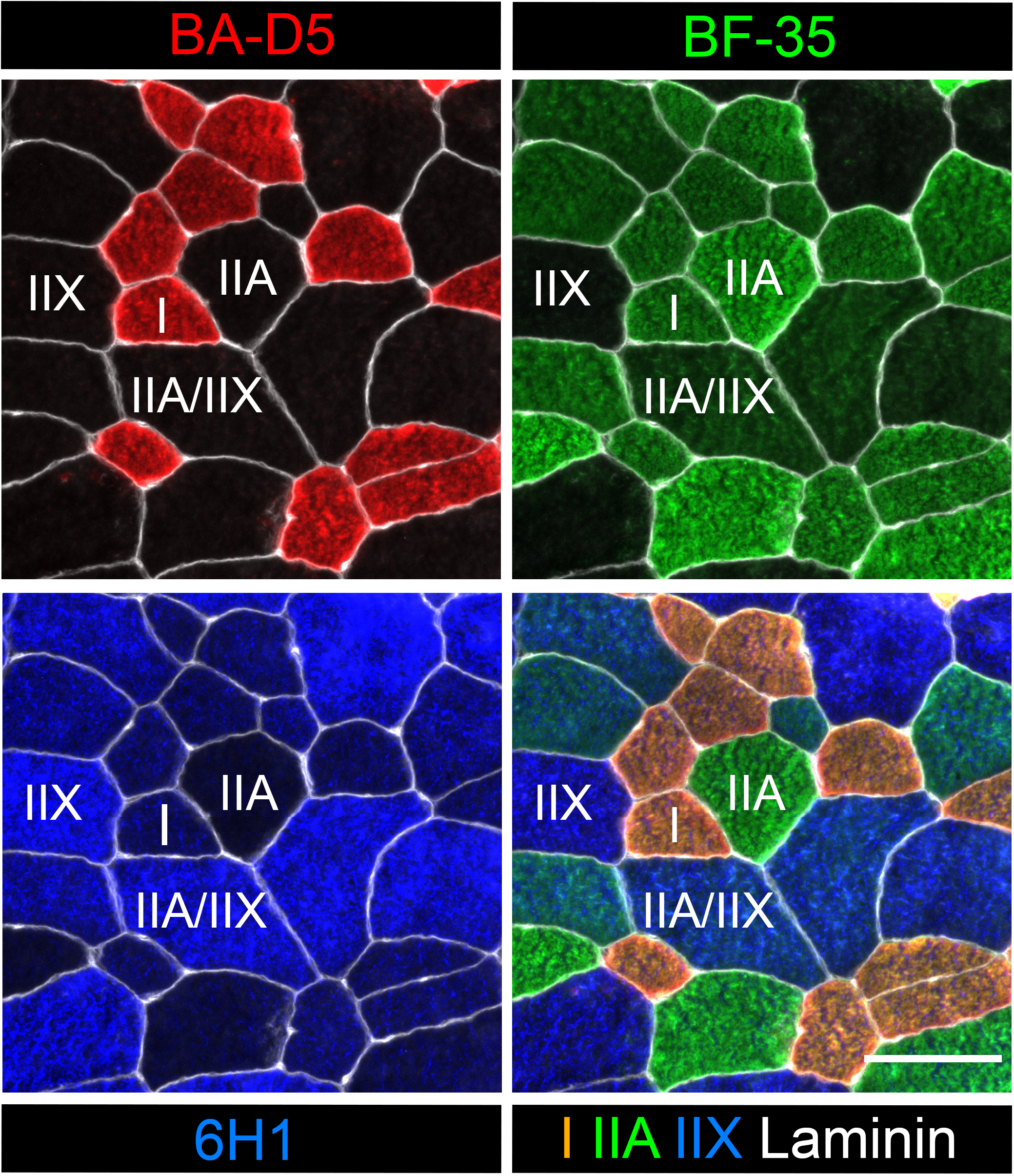
Simultaneous identification of type I, IIA, and IIX myofibers in bovine skeletal muscle: Cross-sections (10 µm) of bovine *longissimus dorsi* muscle biopsy samples were stained with a primary antibody against laminin (ab7463, Rabbit, 1:100) to label the myofiber boundaries, together with a cocktail of mouse monoclonal primary antibodies against MyHC type I (BA-D5, MIgG2b), MyHC type IIX (6H1, MIgM), and all MyHC isoforms except type IIX (BF-35, MIgG1). Primary antibody binding was visualized with Alexa Fluor-conjugated secondary antibodies (1:500) including Goat Anti-Rabbit Alexa Fluor 350 (to detect laminin), Goat Anti-Mouse IgG1 Alexa Fluor 488 (to detect BF-35), Goat Anti-Mouse IgG2b Alexa Fluor 555 (to detect BA-D5), and Goat Anti-Mouse IgM Alexa Fluor 647 (to detect 6H1). Laminin, BA-D5, BF-35, and 6H1 staining were pseudo colored white, red, green, and blue, respectively. Scale bars are 100 µm.

### Similar reactivity of MyHC antibodies when applied individually or in a cocktail

To confirm whether the BA-D5, BF-35, and 6H1 primary antibodies behave similarly when applied as a cocktail, serial bovine muscle cross-sections were stained with these primary antibodies either individually or in combination (**Fig. 4**). The reactivity of these primary antibodies was found to be highly similar when applied in a cocktail when compared to incubation with each individual primary antibody on separate serial sections of the same tissue sample (**Fig. 4**). Notably however, the 6H1 primary antibody while clearly staining type IIX (BF-35^neg^) fibers did cross-react with type I myofibers in bovine muscle both when applied in isolation or in a cocktail format (**Fig. 4**).

**Figure 4:**
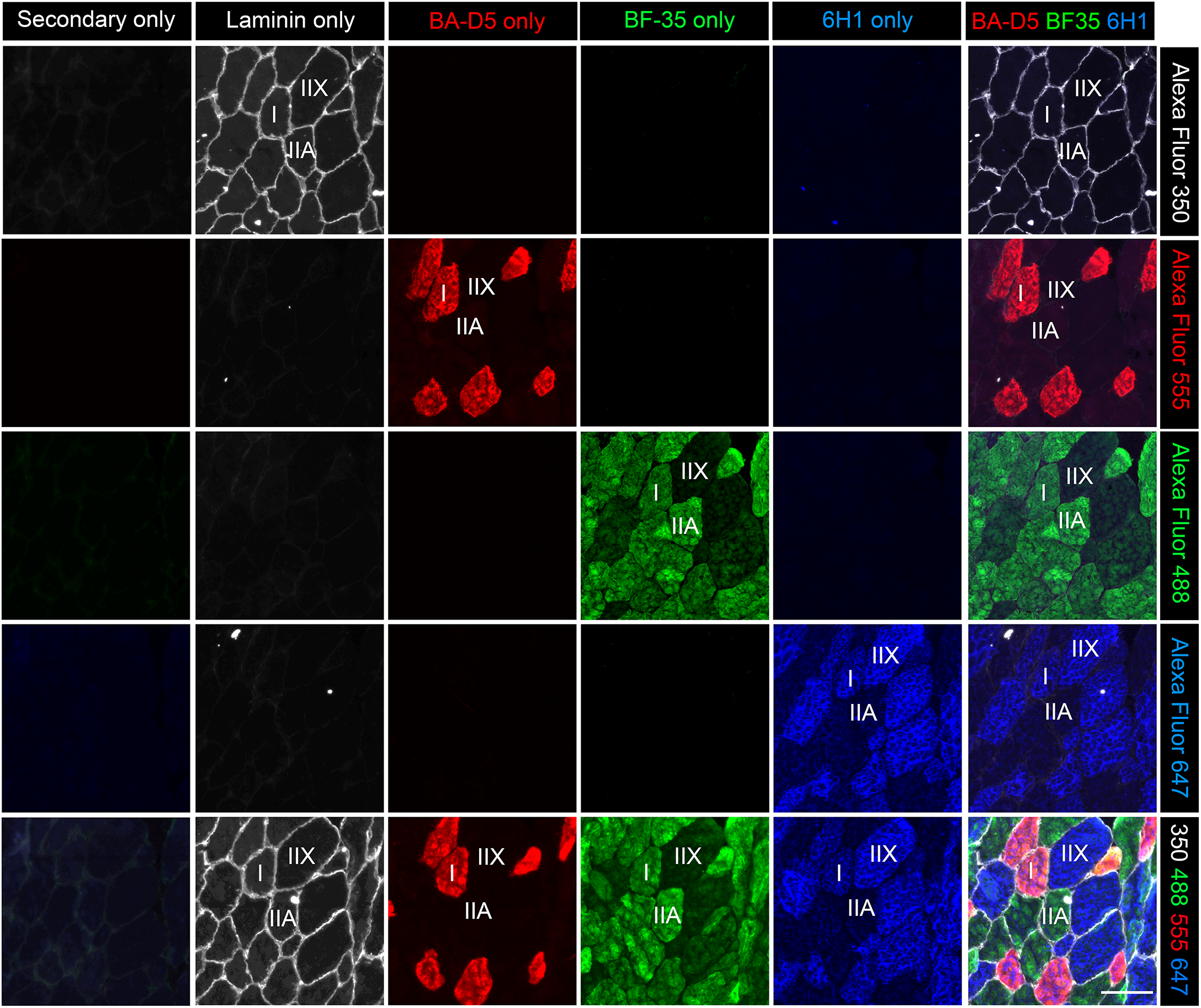
Reactivity of MyHC primary antibodies with bovine skeletal muscle when applied individually or as a cocktail: Serial muscle cross-sections (10 µm) of bovine *longissimus dorsi* muscle biopsy samples were stained with a primary antibody against laminin (ab7463, Rabbit, 1:100) to label the myofiber boundaries together with mouse monoclonal primary antibodies against MyHC type I (BA-D5c, MIgG2b, 1:100), MyHC type IIX (6H1s, MIgM, 1:10), and all MyHC isoforms except type IIX (BF-35c, MIgG1, 1:100), applied either individually or in combination as a cocktail. Primary antibody staining was visualized with a cocktail of Alexa Fluor-conjugated secondary antibodies (1:500) including Goat Anti-Rabbit 350 (to detect laminin), Goat Anti-Mouse IgG2b Alexa Fluor 555 (to detect BA-D5), Goat Anti-Mouse IgM Alexa Fluor 647 (to detect 6H1), and Goat Anti-Mouse IgG1 Alexa Fluor 488 (to detect BF-35). Laminin, BA-D5, BF-35, and 6H1 were pseudo colored white, red, green, and blue, respectively. Scale bars are 100 µm.

### High-throughput automated image analysis of bovine muscle cross-sections

We next questioned whether existing published high-content image analysis methods designed and validated in mice might be successfully applied to the analysis of bovine muscle samples. We focused our initial efforts on the MuscleJ 1.0.2 plugin for FIJI/ImageJ (45). Based on our initial qualitative observations MuscleJ could accurately segment muscle fibers on cross-sections of bovine muscle biopsies when primary antibodies against either laminin or dystrophin were used to stain the myofiber boundaries (**Fig. 5**). MuscleJ could also identify type I (BA-D5^pos^) fibers in bovine skeletal muscle cross-sections although in our hands an adjustment in the default sensitivity threshold for fiber type detection from 3 × standard deviations (Std) from the mean to 1 × Std was required to achieve type I fiber counts that were consistent with manual identification (**Fig. 6**). Therefore, the relative proportions and fiber type-specific CSA of type I fibers (e.g., BA-D5^pos^ cells) vs. type II fibers (e.g., BA-D5^neg^ cells) can be quantified with the default MuscleJ 1.0.2 plugin with some relatively minor modifications to the source code. Unfortunately, however, IIA vs. IIX myofibers cannot be easily distinguished with MuscleJ due to the differences in the reactivity of the SC-71 antibody between rodent and bovine muscle tissue.

**Figure 5:**
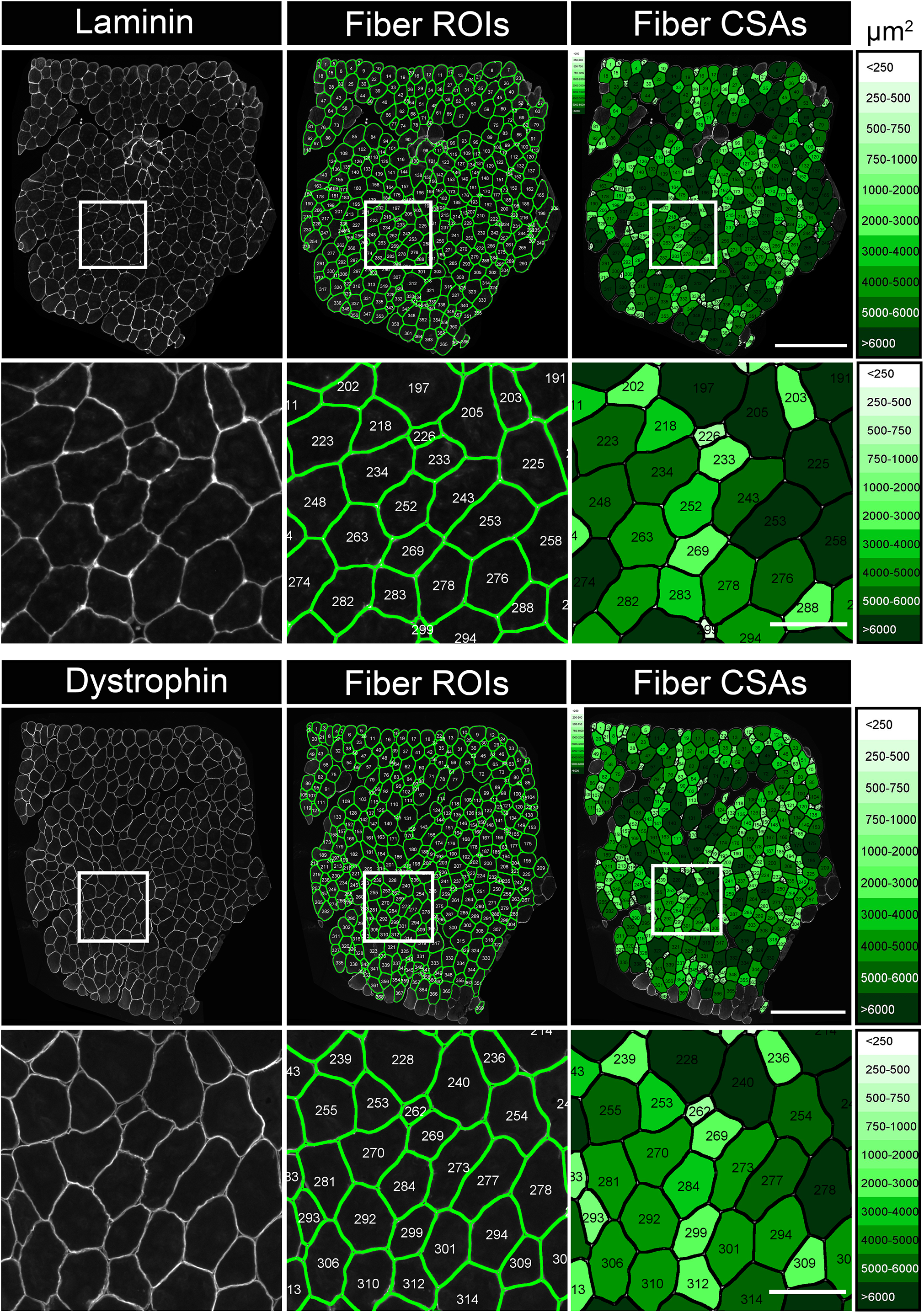
Automated myofiber segmentation on bovine skeletal muscle samples using MuscleJ software: Cross-sections (10 µm) of bovine *longissimus dorsi* muscle biopsy samples were stained with primary antibodies against either laminin (ab7463, Rabbit 1:100) or dystrophin [MANDYS1(3B7)s, MIgG2a, 1:10] to delineate the myofiber boundaries. Primary antibody binding was visualized using Alexa Fluor-conjugated secondary antibodies (1:500) including either Goat Anti-Rabbit Alexa Fluor 350 (to detect laminin) or Goat Anti-Mouse IgG2a Alexa Fluor 647 (to detect dystrophin). Laminin and dystrophin staining was pseudo colored white. Stitched panoramic images of the entire muscle biopsy cross-section were obtained using an automated fluorescent microscope (Echo Revolution) and analyzed using the MuscleJ plugin for FIJI/ImageJ. Segmented myofiber regions of interest (ROIs) were overlaid on the original images and corresponding cartography maps of segmented muscle fibers colored according to their measured cross-sectional areas (CSA) are shown. Scale bars are 400 µm on stitched mosaic images and 100 µm on representative fields of view.

**Figure 6:**
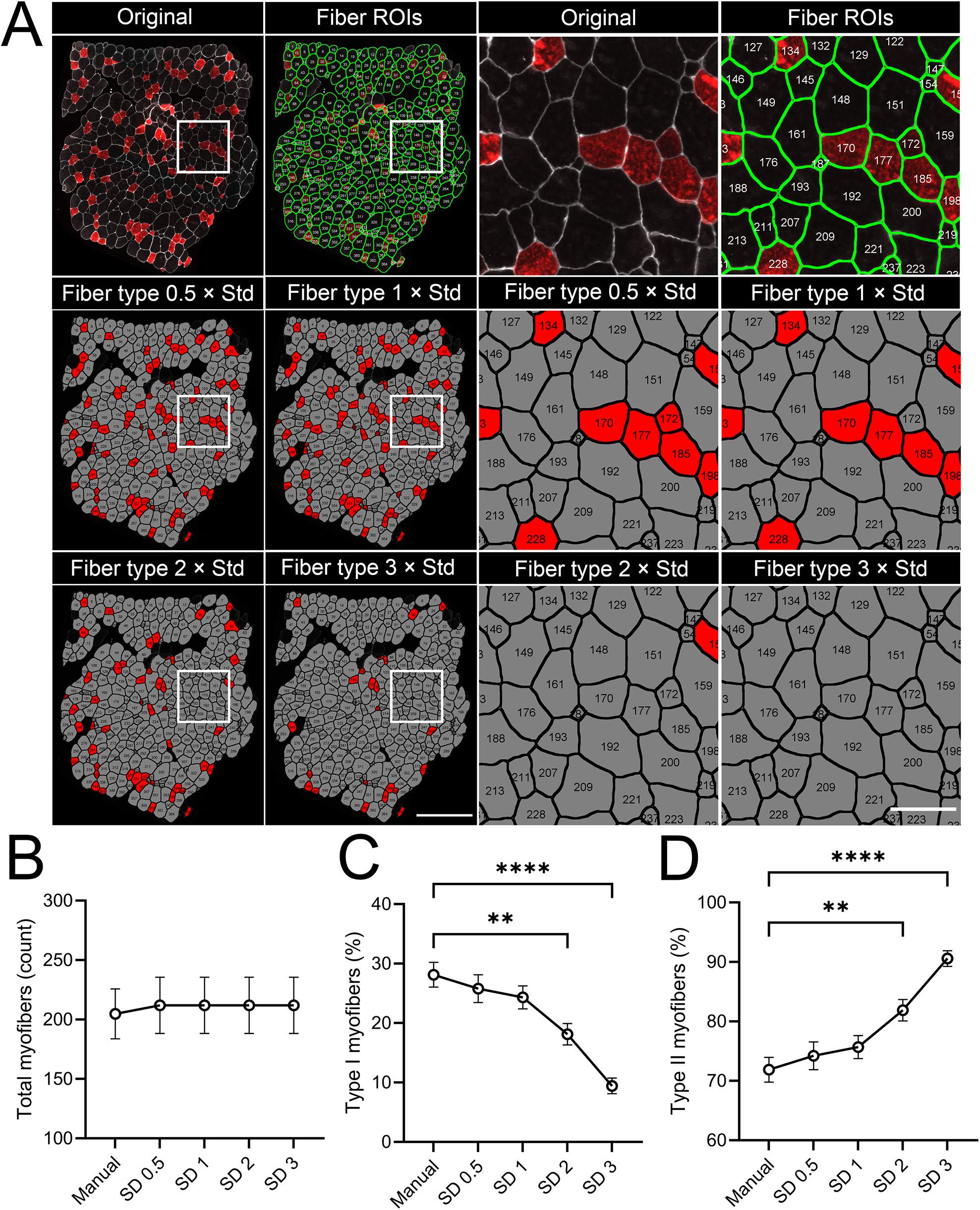
Automated detection of type I myofibers in bovine skeletal muscle using MuscleJ software: **A:** Cross-sections (10 µm) of bovine *longissimus dorsi* muscle biopsy samples were stained with a primary antibody against laminin (ab7463, Rabbit, 1:100) to label the myofiber boundaries in combination with a mouse monoclonal primary antibody against MyHC type I (BA-D5c, MIgG2b, 1:100). Primary antibody binding was visualized using Alexa Fluor-conjugated secondary antibodies (1:500) including Goat Anti-Rabbit Alexa Fluor 647 (to detect laminin) and Goat Anti-Mouse IgG2b Alexa Fluor 555 (to detect BA-D5). Laminin and BA-D5 staining was pseudo colored white and red, respectively. Stitched panoramic images of entire muscle biopsy cross-sections were obtained using an automated fluorescent microscope (Echo Revolution) and analyzed using the MuscleJ 1.0.2 plugin for FIJI/ImageJ with minor custom modification. Segmented myofiber regions of interest (ROIs) were overlaid on the original images and corresponding cartography maps of type I fibers (red) versus non-type I (e.g., type II) fibers (gray) are shown. Scale bars are 400 µm on stitched mosaic images and 100 µm on representative fields of view. **B:** Comparison of total myofiber count data between manual image analysis and MuscleJ. **C:** Comparison of the percentage (%) composition of type I (BA-D5^pos^) fibers between manual image analysis and MuscleJ at different sensitivity thresholds for type I fiber identification. **D:** Comparison of the percentage (%) composition of type II myofibers (BA-D5^neg^) fibers between manual image analysis and MuscleJ at different sensitivity thresholds for type I fiber identification. **B-D:** Data is presented as the mean ± SEM of muscle biopsy samples from n=13 animals. ** and **** denote p<0.01 and p<0.0001 between the indicated groups, respectively.

### Customization of MuscleJ for analysis of bovine muscle fiber type

To attempt to automate the detection and measurement of IIA and IIX myofibers in bovine muscle we next assessed whether MuscleJ could detect staining with the BF-35 primary antibody. While MuscleJ was not originally designed to identify BF-35 staining, we found that the function originally designed to detect type IIB myofibers on mouse muscle samples worked well to identify BF-35^pos^ myofibers on bovine muscle cross-sections (**Fig. 7**). Moreover, when combined with detection of type I fibers (BA-D5^pos^ cells) using a staining intensity threshold of 1 × Std from the mean MuscleJ could also identify type I (BA-D5^pos^ cells), type IIA (BA-D5^neg^BF-35^pos^ cells), and type IIX fibers (BA-D5^neg^BF-35^neg^ cells) on a single tissue section (**Fig. 7**). According to these new designations for bovine muscle fiber type classification we made extensive custom modifications to the MuscleJ 1.0.2 source code, user interface (UI), and results outputs. Since an adjustment in the default sensitivity threshold of type I fiber identification was required in our hands and this may potentially vary between laboratories, we incorporated a new UI option to enable users to easily set custom intensity thresholds for positive fiber type identification. We also added new UI options to generate cartography maps with custom color schemes. Finally, we added the option to generate multiple different cartography files e.g., myofiber area and fiber type simultaneously from a single run. We named this new custom bovine-specific MuscleJ 1.0.2 version MuscleBos after the scientific bovine genus *Bos* (from Latin bōs: cow, ox, bull).

**Figure 7:**
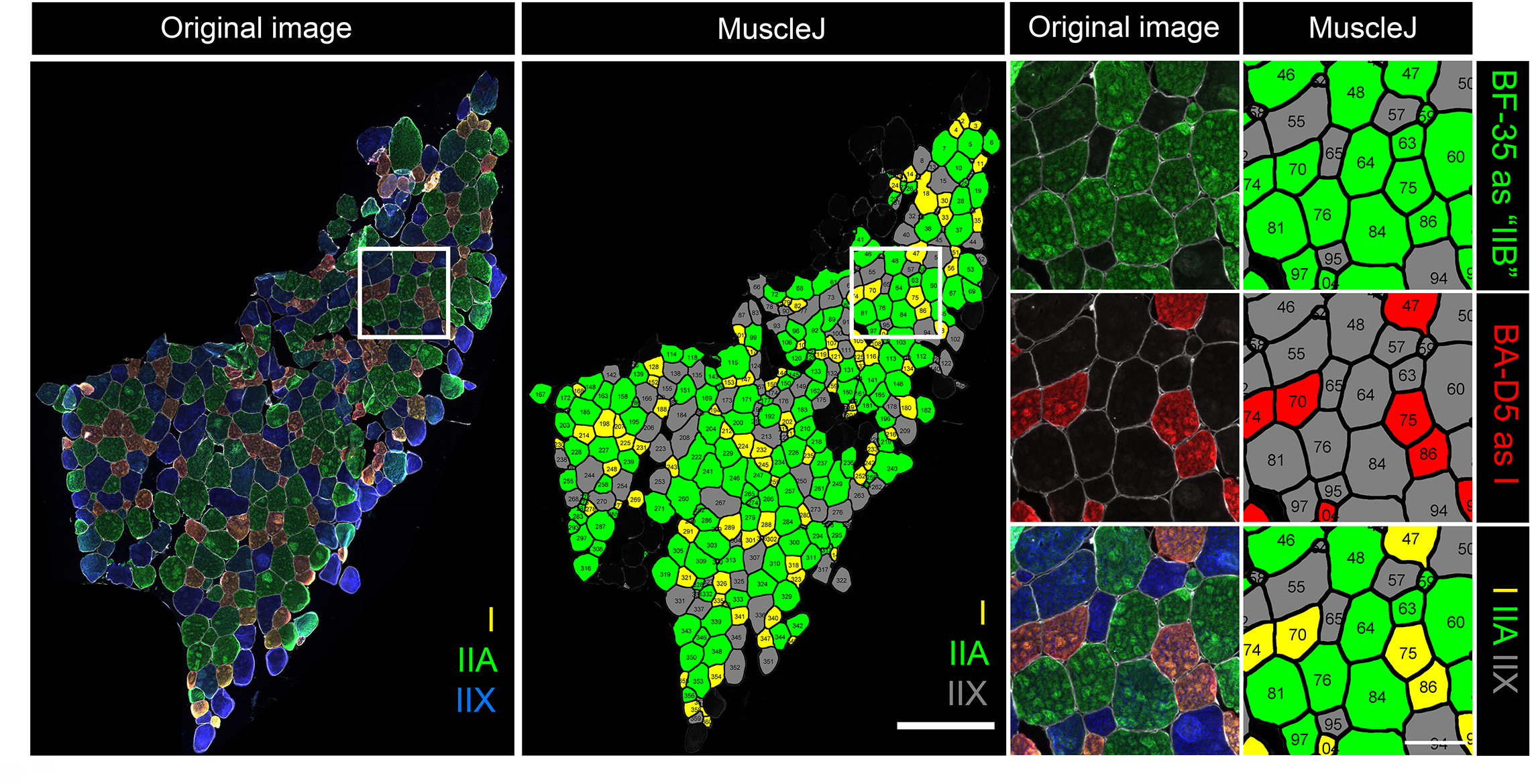
Simultaneous detection of type I, IIA, and IIX myofibers in bovine skeletal muscle using customized MuscleJ software: Bovine *longissimus dorsi* muscle biopsy cross-sections were stained with a primary antibody against laminin (ab7463, Rabbit, 1:100) to label the myofiber boundaries together with a cocktail of mouse monoclonal primary antibodies against MyHC type I (BA-D5c, MIgG2b, 1:100), all but IIX (BF-35c, MIgG1, 1:100), and type IIX (6H1s, IgM, 1:10). Primary antibody binding was detected using Alexa Fluor-conjugated secondary antibodies (1:500) including Goat Anti-Rabbit Alexa Fluor 350 (to detect laminin), Goat Anti-Mouse IgG2b Alexa Fluor 555 (to detect BA-D5), Goat Anti-Mouse IgG1 Alexa Fluor 488 (to detect BF-35), and Goat Anti-Mouse IgM Alexa Fluor 647 (to detect 6H1). Laminin, BA-D5, SC-71, and 6H1 staining was pseudo colored white, red, green, and blue, respectively. Stitched panoramic images of the entire muscle biopsy cross-section were obtained using an automated fluorescent microscope (Echo Revolution) and analyzed using the MuscleJ plugin for FIJI/ImageJ with custom modifications. Original images and fiber type cartography maps of type I (red), type IIA (green), and type IIX (gray) are shown. Scale bars are 400 µm on stitched mosaic images and 100 µm on representative fields of view.

### Validation of MuscleBos software

We next sought to validate our initial prototype version of MuscleBos via side-by-side comparisons with manual image analysis. A trained expert manually analyzed a random selection of images of *longissimus dorsi* biopsy cross-sections obtained from multiparous Holstein dairy cows (58). The same images were analyzed in parallel using MuscleBos software (**Fig. 8A**). Manual analysis identified a total of 2755 fibers while MuscleBos counted a total of 2661 fibers. The average number of fibers per image identified by MuscleBos was 211.92 which did not differ significantly from manual fiber counts of 204.69 (**Fig. 8B**). MuscleBos completed this task in ∼13 minutes achieving an average speed of ∼200 fiber/min. In contrast, it took the trained expert approximately 13 hours working at a pace of ∼3.5 fiber/min to complete this task. MuscleBos was therefore approximately 60 times faster than manual image analysis.

**Figure 8:**
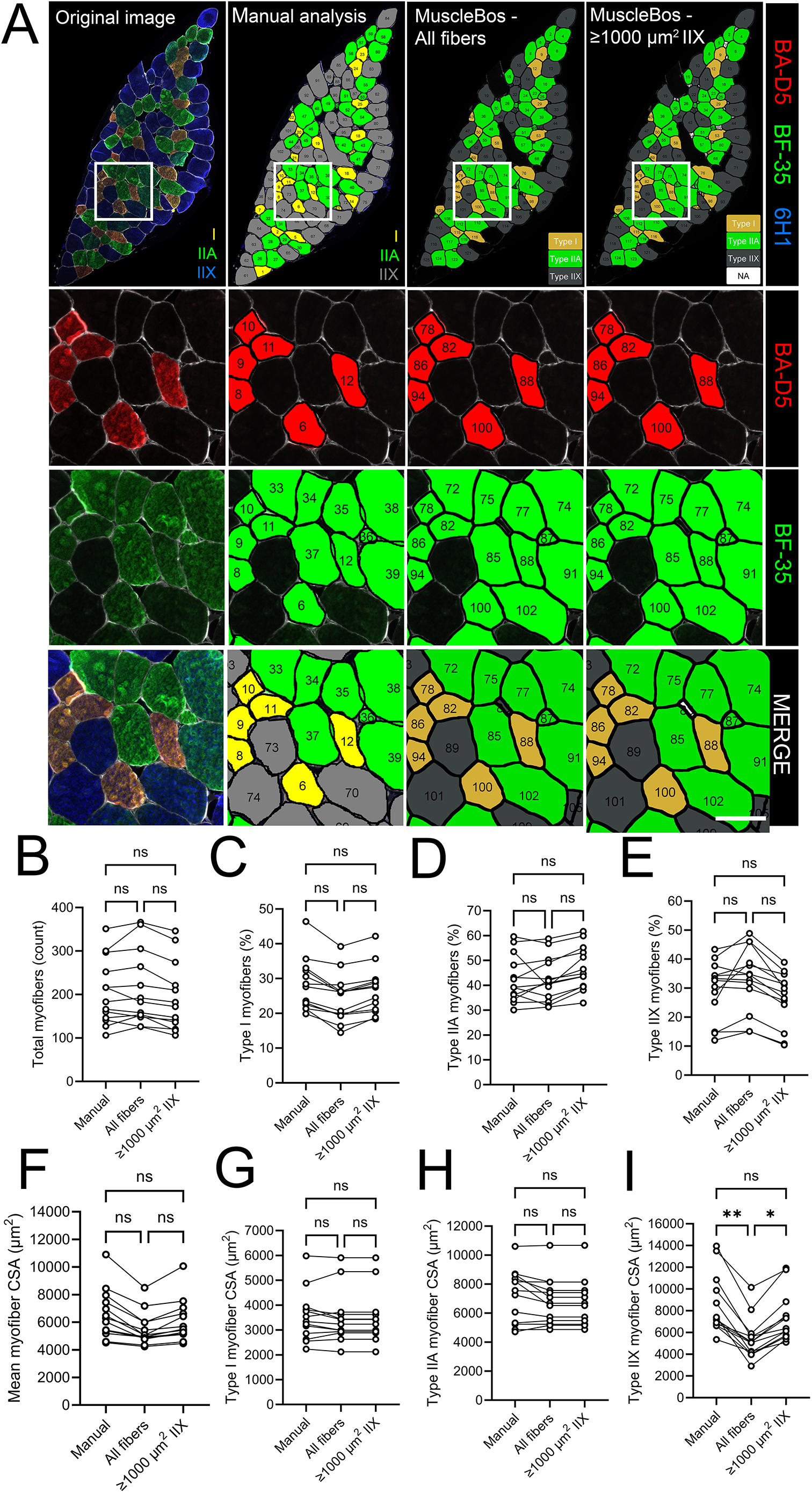
Validation of MuscleBos vs. manual quantification: **A:** Bovine *longissimus dorsi* muscle cross-section (10 µm) were stained with a primary antibody against laminin (ab7463, Rabbit, 1:100) to label the myofiber boundaries together with a cocktail of mouse monoclonal antibodies against MyHC type I (BA-D5c, MIgG2b, 1:100), MyHC type IIX (6H1s, MIgM, 1:10), and all MyHC types except IIX (BF-35c, MIgG1, 1:100). Primary antibody staining was visualized with Alexa Fluor-conjugated secondary antibodies (1:500) including Goat Anti-Rabbit Alexa Fluor 350 (to detect laminin), Goat Anti-Mouse IgG2b Alexa Fluor 555 (to detect BA-D5), Goat Anti-Mouse IgG1 Alexa Fluor 488 (to detect BF-35), and Goat Anti-Mouse IgM Alexa Fluor 647 (to detect 6H1). Laminin, BA-D5, BF-35, and 6H1 were pseudo colored white, red, green, and blue, respectively. Stitched panoramic images of the entire muscle biopsy cross-section were obtained using an automated fluorescent microscope (Echo Revolution) and analyzed using the MuscleBos plugin for FIJI/ImageJ. An original image and corresponding manually or MuscleBos generated fiber type cartography maps are shown. Scale bars on representative fields of view are 100 µm. **B–E:** Comparison of manual and MuscleBos analysis of the same bovine muscle cross-sections for total myofiber count (**B**), the percentage of type I myofibers (**C**), the percentage of type IIA myofibers (**D**), and the percentage of type IIX myofibers (**E**). **F–H:** Comparison of manual and MuscleBos analysis of the same bovine muscle cross-sections for mean myofiber cross-sectional area (CSA) measurements for all fibers irrespective of type (**F**), type I fibers (**G**), type IIA fibers (**H**), and type IIX fibers (**I**). Dot plots represent data from each individual animal (biological replicates) for n=13 cows. NS denotes no significant difference between manual analysis and MuscleBos. * and ** denote p<0.05 and p<0.01 between the indicated groups, respectively.

The percentage of type I (**Fig. 8C**), IIA (**Fig. 8D**), and IIX (**Fig. 8E**) fibers did not differ between MuscleBos and manual image analysis. Total myofiber CSA (**Fig 8F**), type I fiber CSA (**Fig. 8G**) and type IIA fiber CSA (**Fig. 8H**) measurements were also similar between MuscleBos and manual analysis. Our initial version of MuscleBos did measure the CSA of the type IIX myofibers to be significantly smaller when compared with manual image analysis (**Fig. 8I**). This was found to be due to MuscleBos identifying some small unstained spaces between adjacent myofibers as type IIX myofibers (BA-D5^neg^BF-35^neg^ cells) (**Fig 8A**). To address this potential limitation, we incorporated into the MuscleBos UI a new option to set user defined fiber type specific CSA exclusion thresholds. For type IIX fibers we set a default exclusion threshold of ≤1000 µm^2^ since very few, if any, type IIX myofibers below this size were identified in manual image analysis. Excluding these very small unstained objects did not influence the average myofiber count (**Fig. 8B**), the percentage of type I (**Fig. 8C**), type IIA (**Fig. 8D**), and IIX (**Fig. 8E**) fibers, or the total mean myofiber CSA (**Fig. 8F**). Nevertheless, applying this option improved the consistency between MuscleBos and manual analysis for measurement of the mean type IIX fiber CSA (**Fig. 8I**).

### Application of MuscleBos software

We next applied MuscleBos to the analysis of muscle biopsy samples obtained from a previously published study (58). In this prior study, *longissimus dorsi* muscle depth was measured in multiparous Holstein dairy cows via ultrasonography at 42-days before expected calving and cows were assigned based on baseline muscle depth to either the high muscle group (HM, ≥4.6 cm, n=26) or low muscle group (LM, ≤4.6 cm n=22). Cows were further randomized to receive dietary supplementation with control (CON) or branched-chain volatile fatty acid (BCVFA) supplementation from 42 days before expected calving until parturition. Biopsy samples were obtained from the *longissimus dorsi* muscle at 21-days before expected calving and at 21-days postpartum. Dietary supplementation with BCVFAs did not influence any indices of muscle fiber size or type. Therefore, data is presented here pooled over dietary supplementation treatment.

MuscleBos completed fiber type analysis of a total of 27,508 fibers from available immunofluorescent images of a total of 90 biopsy samples in a duration of 2 hours and 17 min achieving an average rate of ∼200 fibers/min. Representative fiber type staining of cross-sections *longissimus dorsi* muscle biopsy samples from HM and LM cows obtained pre and postpartum are shown in **Fig. 9A**. The overall fiber type profile of these samples was found to be 27.27% type I, 47.33% type IIA, and 25.41% type IIX (**Fig. 9B**). The mean CSA of type I myofibers was lower than both type IIA and type IIX fibers (**Fig. 9C**). There was a significant effect of both group and time, but no group × time interaction effect for myofiber CSA (**Fig. 9D**). HM cows showed a larger fiber CSA than LM cows and fiber CSA decreased from pre to postpartum (**Fig. 9D**). There was no significant effect of group, time, or group × time interaction effect for type I fiber CSA (**Fig. 9E**). Type IIA fiber CSA showed a significant main effect of both group and time, but no group × time interaction (**Fig. 9F)**. Type IIA myofiber CSA was larger in HM vs. LM cows and decreased from pre- to post-parturition (**Fig. 9F)**. Type IIX myofiber CSA showed a significant main effect of group, but no effect of time, or group × time interaction effect (**Fig. 9G)**. Type IIX fiber CSA was overall higher in HM vs LM cows (**Fig. 9G)**. There was no significant effect of time, group, or time by group interaction for the percentage of type I myofibers (**Fig. 9H**). The percentage of type IIA myofibers showed a significant main effect of group and a statistical trend for a group by time interaction effect (**Fig. 9I**). HM cows had a greater proportion of type IIA fibers than LM cows and this was especially true in the postpartum biopsy samples (**Fig. 9I**). The proportion of type IIX myofibers showed a significant main effect of group and a group × time interaction effect (**Fig. 9J**). HM cows had a lower percentage of type IIX myofibers in the postpartum period when compared to LM cows (**Fig. 9J**).

**Figure 9:**
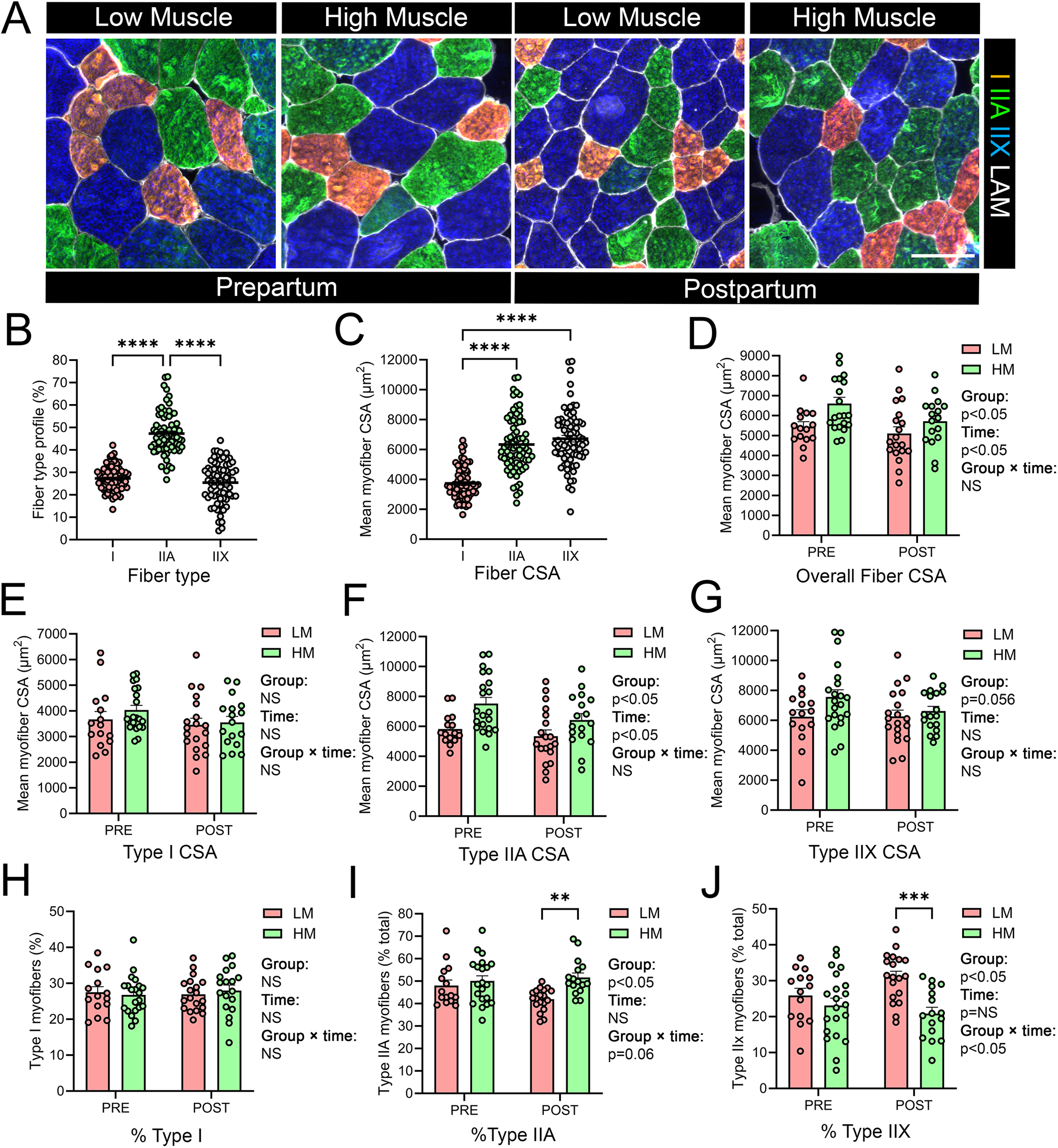
Application of MuscleBos to the characterization of bovine muscle phenotype: **(A):** Muscle biopsy samples were obtained from the *longissimus dorsi* of high muscle (HM) or low muscle (LM) multiparous dairy cows at 21-days before expected calving (PRE) and at 21-days postpartum (POST). Biopsy cross-sections (10 µm) were stained with a primary antibody against laminin (ab7463, Rabbit, 1:100) to label the myofiber boundaries together with a cocktail of mouse monoclonal antibodies against MyHC type I (BA-D5c, MIgG2b, 1:100), MyHC type IIX (6H1s, MIgM, 1:10), and all MyHC types except IIX (BF-35c, MIgG1, 1:10). Primary antibody staining was visualized with Alexa Fluor conjugated secondary antibodies (1:500) including Goat Anti-Rabbit Alexa Fluor 350 (to detect laminin), Goat Anti-Mouse IgG2b Alexa Fluor 555 (to detect BA-D5), Goat Anti-Mouse IgG1 Alexa Fluor 488 (to detect BF-35), and Goat Anti-Mouse IgM Alexa Fluor 647 (to detect 6H1). Laminin, BA-D5, BF-35, and 6H1 were pseudo colored white, red, green, and blue, respectively. Scale bars are 100 µm. **B-J:** Stitched panoramic images of the entire muscle biopsy cross-section were obtained using an automated fluorescent microscope (Echo Revolution) and analyzed using the MuscleBos plugin for FIJI/ImageJ. **B:** The overall fiber type profile of pooled bovine *longissimus dorsi* muscle biopsy samples showing the percentage composition of type I, type IIA, and type IIX fibers as determined by MuscleBos. **C:** The mean cross-sectional area (CSA) type I, type IIA, and type IIX myofibers in pooled bovine *longissimus dorsi* muscle biopsy samples as determined by MuscleBos. **D:** Comparison of mean myofiber CSA irrespective of fiber type between bovine *longissimus dorsi* muscle biopsy samples obtained from HM or LM cows at PRE and POST parturition. **E-G**: Comparison of the mean CSA of type I **(E)**, type IIA **(F)**, and type IIX **(G)** myofibers in bovine *longissimus dorsi* muscle biopsy samples obtained from HM or LM cows at PRE and POST parturition **H-J**: Comparison of percentage fiber type composition of type I **(H)**, type IIA **(I)**, and type IIX **(J)** myofibers in bovine *longissimus dorsi* muscle biopsy samples obtained from HM or LM cows PRE and POST parturition. Values are mean ± SEM with dot plots representing data obtained from each individual animal (biological replicates). Symbols denote *p<0.05, **p<0.01, and ***p<0.001, and ****p<0.0001 between the indicated groups.

## Discussion

The present study established and validated a high-throughput immunofluorescence-based method for characterizing bovine skeletal muscle fiber type composition and fiber-type specific myofiber size. By combining optimized antibody cocktails with automated fluorescent microscopy, and high-content image analysis using the new MuscleBos plugin for FIJI/ImageJ, we demonstrated that reliable rapid automatic identification and quantification of type I, IIA, and IIX myofibers can be achieved on bovine muscle cross-sections. This approach substantially reduces the time and effort required for image analysis while maintaining accuracy comparable to manual annotation, enabling its application in large-scale studies of bovine muscle biology and meat science.

Early studies employing traditional histochemical myosin ATPase staining techniques reported that bovine skeletal muscle contained three muscle fiber types including I, IIA, and IIB (64). However, consistent with our findings more recent studies using fiber type specific monoclonal primary antibodies have reported that MyHC IIX is the only fast-twitch glycolytic isoform that is present in trunk and limb muscles of bovine species (50, 65–67). The presence of type IIB muscle fibers therefore generally appears restricted to small mammals such as mice and rats. Interestingly, the MyHC IIB isoform is however widely expressed in swine muscle (68). While apparently lacking in bovine limb and trunk muscles, type IIB fibers have also been detected in specialized eye muscles of cattle (65, 67). Some studies have further suggested that the mRNA encoding MyHC type IIB (*Myh4*) is indeed expressed together with variably detectable protein levels in limb muscles of some specialized breeds of cattle such as the French beef breed, Blonde d’Aquitaine (69–71). On this basis, our method may not be applicable to all bovine muscle types or cattle breeds. Nevertheless, the methods described here should be generally applicable to analysis of limb and trunk muscles of the most common breeds of dairy and beef cattle.

Another key finding of our study is that the widely used SC-71 primary antibody, which in rodents reacts selectively with type IIA fibers, exhibited broader reactivity in bovine muscle. Specifically, in the bovine *longissimus dorsi* muscle biopsy samples analyzed here, the SC-71 antibody clearly labeled all type II myofibers, including both IIA and IIX fibers. Although initially unexpected based on our prior experience, this finding is indeed consistent with several previously published reports in cattle (15, 49, 50, 67, 72). In human skeletal muscle, the SC-71 antibody also cross-reacts with IIX muscle fibers, albeit with notably dimmer staining than type IIA fibers, thereby potentially allowing for type IIA and IIX fibers to be visually distinguished based on relative staining intensity (73). In contrast, we observed no such difference in the staining intensity between type IIA and IIX myofiber populations in bovine muscle. This observation precludes use of the SC-71 primary antibody to distinguish type IIA and IIX myofibers in bovine skeletal muscle. This species-specific limitation underscores the necessity of robust primary antibody validation in non-rodent samples and highlights the importance of establishing alternative unique antibody combinations to accurately resolve muscle fiber types in cattle.

We demonstrate that a cocktail of BA-D5 (type I-specific), BF-35 (all but IIX-specific), and 6H1 (IIX-specific) antibodies provide robust differentiation of the three major adult bovine fiber types on a single tissue section. This was expected based on early reports which already comprehensively characterized the reactivity of each of these primary antibodies in isolation on bovine muscle (15, 65, 67, 72). Indeed, similar primary antibody cocktails as described here have been successfully used by several other groups to enable simultaneously detection of the three major adult bovine MyHC isoforms (I, IIA, and IIX) in recent years (28, 29, 33, 49–52, 54–56). Importantly, when tested both individually and in combination, we found that BA-D5, BF-35, and 6H1 primary antibodies displayed consistent reactivity, supporting their utility in multiplexed staining protocols. Nevertheless, some cross-reactivity of the type IIX specific 6H1 primary antibody with type I fibers was noted in our hands. This finding is consistent a prior study by Song et al. 2020 who also reported some cross-reactivity of the 6H1 antibody with type I muscle fibers in beef cattle (50). Therefore, it appears preferable to characterize bovine type IIX myofibers based on their relative lack of BF-35 signal, either alone or in combination with their positive reactivity with 6H1, rather than based on 6H1 staining alone.

We found that the default version of MuscleJ 1.0.2 which was originally designed and validated for use on mouse muscle samples could accurately segment myofiber boundaries on bovine muscle biopsy cross-sections stained with either laminin or dystrophin. Moreover, type I fibers could readily be distinguished from non-type I (e.g., pooled type IIA + IIX) fibers. However, on the samples analyzed here adjustment of the sensitivity threshold was required to accurately capture all BA-D5⁺ fibers that were counted in manual image analysis, emphasizing the potential need for optimization of software parameters for individual laboratory-, species-, and/or antibody-specific contexts. When BF-35 and 6H1 primary antibodies were incorporated into a cocktail together with BA-D5, our new customized bovine-specific version of MuscleJ 1.0.2 that we termed MuscleBos could also successfully automate classification of type I, IIA, and IIX fibers, providing powerful quantitative measures of complete bovine muscle fiber type distribution and fiber type–specific CSA.

Validation against expert manual annotation confirmed that MuscleBos delivered highly comparable results. Our initial prototype version of MuscleBos produced similar total fiber counts, fiber type proportions, and myofiber CSA values when compared to manual image analysis. However, our initial version that was based on the default MuscleJ myofiber segmentation code did consistently underestimate type IIX myofiber CSA on our bovine muscle samples. MuscleJ was originally designed to identify type IIX myofibers based on a lack of staining with other primary antibodies. Because of this it sometimes incorrectly identified spaces between adjacent muscle fibers as small type IIX fibers on the samples tested here. This limitation may reflect the larger average size of bovine vs. rodent muscle fibers and/or the looser packing of muscle fibers in muscle biopsy samples from larger species when compared to complete cross-sections of entire mouse muscle samples in which myofibers are more densely organized. To address this potential limitation, we incorporated into MuscleBos the ability to add user defined fiber type specific CSA exclusion thresholds. Excluding unstained objects (e.g., IIX fibers) with CSA measurements of ≤1000 µm^2^ further improved the consistency between MuscleBos and manual image analysis.

When applied to biopsy samples obtained from dairy cows differing in prepartum *longissimus dorsi* muscle depth, MuscleBos revealed statistically significant differences in both percentage muscle fiber type composition and fiber type specific myofiber CSA. Specifically, the HM cows displayed larger CSA of type IIA and IIX (but not type I) myofibers and had a relatively higher proportion of type IIA fibers during the post-partum period. In contrast, LM cows showed lower type IIA and IIX (but not type I) myofiber CSA and a greater proportion of type IIX fibers during the postpartum period. We were further able to demonstrate fiber type-specific atrophy of type IIA myofibers from pre- to post-partum. These data show for the first time that fast oxidative-glycolytic fibers are preferentially mobilized during the transition period during which dairy cows lose large amounts of skeletal muscle mass. Overall, these findings demonstrate that automated image analysis can capture potentially physiologically relevant variation changes in muscle fiber morphology and can serve as a powerful tool to study the underlying cellular basis of skeletal muscle remodeling in response to adaptation to lactation and/or nutritional changes in dairy cattle.

We based MuscleBos on the MuscleJ 1.0.2 plugin for FIJI/ImageJ (45). This decision was primarily due to our prior familiarity with MuscleJ and recent success in utilizing this plugin for the automated quantitative analysis of fiber type on rodent skeletal muscle samples (60, 61, 63, 74). Nevertheless, the immunofluorescent staining and imaging protocol described here should, in theory, also be compatible with several other freely available muscle specific image analysis software options. The first prerequisite would be the ability to identify staining of various MyHC-specific primary antibodies following segmentation of the myofiber boundaries. This is a feature that is offered by most (35–38, 40–45, 47), but not all software options (39, 46, 57). The second requirement would be the ability to identify myofibers that are double positive for two different primary antibodies e.g., type IIA fibers as BA-D5^pos^/BF-35^pos^ cells. Several other software originally designed and validated on mouse samples do allow for the potential detection of hybrid muscle fiber types expressing two MyHC types, most commonly I-IIA myofibers (e.g., BA-D5^pos^/SC-71^pos^ cells) (35, 36, 38, 40–44). Therefore, in theory the staining and imaging methods established here may also be compatible with QuantiMus (35), SMASH (36), MyoSight (38), Myosoft (40), the unnamed ImageJ plugin of Bergmeister et al. 2016 (41, 42), Cellpose + LabelsToRois (43), and MuscleJ2 (44). However, some degree of customization, optimization, and validation would most likely be required, especially for those that only allow for fully automated analysis without any user input or customization (e.g., 44, 47). On the other hand, the bovine-specific primary antibody cocktails used here would most likely be incompatible with MyoVision (37), Open-CSAM (39), MyoView (47), the unnamed FIJI/ImageJ plugin by Reyes-Fernandez (48), MyoV (57), and MyoAnalyst (46).

The methodology described here has several limitations. Firstly, the combination of BA-D5, BF-35, and 6H1 primary antibodies does not allow for potential detection of hybrid type I-IIA myofibers. Hybrid type I-IIA fibers are rarely observed in bovine muscle (50). Nevertheless, they could potentially be identified via manual analysis of serial sections stained with a cocktail of BA-D5 and SC-71 (**Fig. 1 & 2**). The most commonly observed hybrid fiber type present in bovine muscle are IIA-IIX fibers (50). Hybrid IIA-IIX myofibers can potentially be identified as BA-D5^neg^/BF-35^pos^/6H1^pos^ cells with the cocktail of BA-D5, BF-35, and 6H1 primary antibodies used here (**Fig. 4**). Nevertheless, we chose not to include automated identification of this hybrid fiber type category in MuscleBos, in part, due to the apparent cross-reactivity of the 6H1 primary antibody observed with type I myofibers in bovine muscle. Finally, the accuracy of both fiber type and CSA determination with MuscleBos, like MuscleJ 1.0.2 on which it was based, is dependent on the quality of staining antibody staining of the myofiber boundaries e.g., with laminin, dystrophin, or wheat germ agglutinin. Therefore, dim or uneven staining of the myofiber boundaries may lead to inaccurate fiber segmentation results which would compromise the downstream accuracy of both fiber type proportion and myofiber CSA measurements.

## Conclusions

In conclusion, this study provides a validated framework for rapid, high-throughput analysis of bovine muscle fiber type and morphology incorporating automated fluorescence microscopy and automated image analysis with MuscleBos that is accurate, efficient, and scalable. The methodological advances described here should facilitate more comprehensive investigations into bovine muscle biology, with implications for animal physiology, performance, and meat quality.

## Declarations

### Ethics approval and consent to participate

All animal procedures were approved by the Purdue University Institutional Animal Care and Use Committee (IACUC) (protocol #2109002197).

### Consent for publication

Not applicable.

### Availability of data and materials

All data generated or analyzed during this study are included in this published article and its supplementary information files.

### Authors’ contributions

HR performed wet laboratory analysis including tissue cryosectioning, immunofluorescent staining, and fluorescent microscopy. HR also edited the MuscleJ 1.0.2 source code to create the first prototype version of MuscleBos, contributed to method development, and wrote the initial draft of the manuscript. KMG performed animal studies from which biological samples were obtained and collected all bovine muscle biopsy samples. RKC performed all manual image analysis against which MuscleBos was validated. JPB acquired funding, supervised trainees (KMG and RKC), and oversaw the animal studies from which muscle biopsy samples were obtained. JAP worked closely with HR and JFM in method development and wrote the final version of the MuscleBos plugin for FIJI/ImageJ. JFM conceived the studies, acquired funding, supervised trainees (HR), worked with HR and JAP in the creation of MuscleBos, and worked with HR to write and edit the final version of the manuscript. All authors read and approved of the final manuscript.

## Supporting information

MuscleBos User Guide

MuscleBos V1-0

## Acknowledgments

We would like to thank S. Haag, M. Simonds and the Purdue Dairy Farm staff for their assistance with this project. Monoclonal antibodies including MANDYS1(3B7)s, BA-D5c, SC-71c, 6H1s, and BF-35c were obtained from the Developmental Studies Hybridoma Bank (DSHB), created by the NICHD of the NIH and maintained at The University of Iowa, Department of Biology, Iowa City, IA 52242.

