## Supplementary material for "Rapid High-Throughput Analysis of Bovine Skeletal Muscle Fiber Morphology via Automated Fluorescent Microscopy and MuscleBos software": MuscleBos User Guide

**MuscleBos V1-0 User Tutorial**

**Step 1: Preparation of images for MuscleBos analysis**

- MuscleBos utilizes the “*Original File Format”* Data Format option originally available in MuscleJ 1.0.2.
    - The “*TIFF (16bits) by channel”* Data Format option that was originally available in MuscleJ 1.0.2 has been removed from the MuscleBos plugin.

- Like MuscleJ 1.0.2 on which it was based, MuscleBos is compatible with “*Original File Format*” image files readable via the *“Bioformat Importer Plugin”* for FIJI/ImageJ such as czi, lsm, lif, nd2 etc.

- Merged composite TIFF (16-bit) images files that contain all fluorescent channels (e.g., a stack file from FIJI/ImageJ) are also compatible with MuscleBos.
    - If the image(s) to analyzed are in single channel TIFF (16-bit) format, then first merge them in FIJI/ImageJ prior to MuscleBos analysis to obtain a corresponding multichannel composite 16-bit TIFF image.
        - Open the individual single channel 16-bit TIFF images to be analyzed in FIJI/ImageJ.
            - Select: “*Image*” → “*Color*” → “*Merge Channels…*”

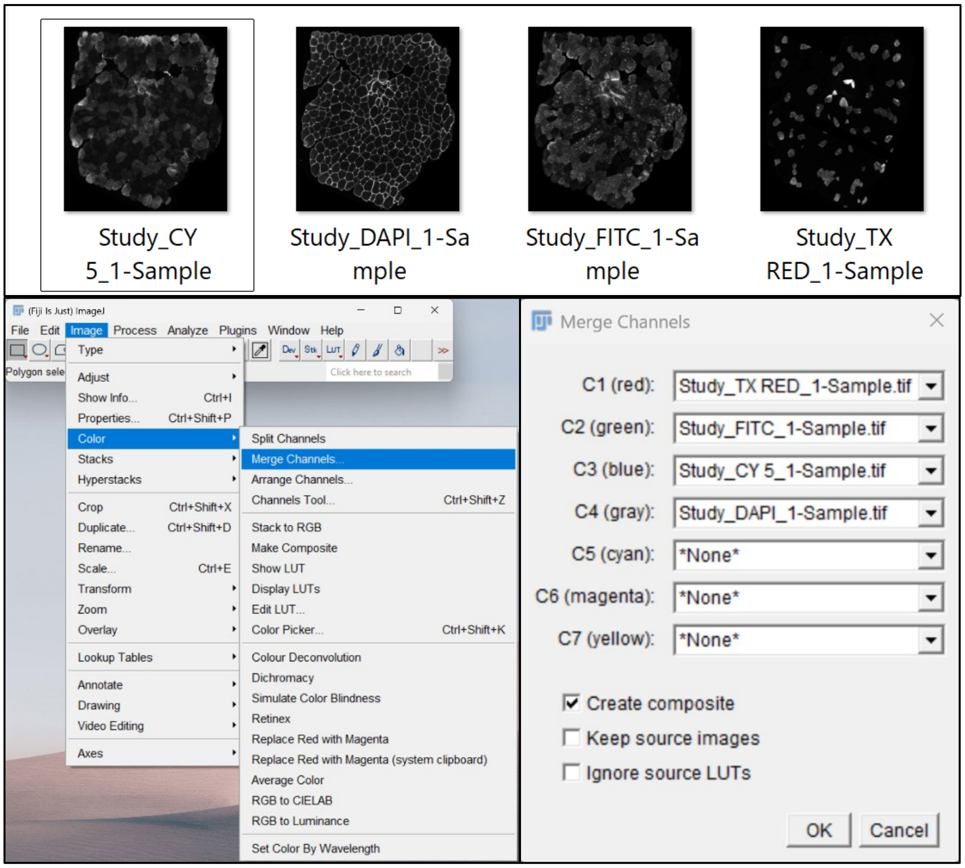

        - Assign each of the individual single channel 16-bit TIFF images either their true emission color or the desired pseudocolor.
            - C1 (red)
            - C2(green)
            - C3 (blue
            - C4 (gray)
            - C5 (cyan)
            - C6 (magenta)
            - C7 (yellow)
        - Select "*Create composite*"
            - Select "*OK*".

    - For composite multichannel 16-bit TIFF images merged as described above the fluorecent channels are ordered: C1 (red), C2 (green), C3 (blue), C4 (gray), C5 (cyan), C6 (magenta), C7 (yellow).
        - If a given color is either not assigned the remaining channels are named in order:
            - C1 (green), C2 (blue), C3 (gray) for a 3-color composite image lacking a red channel.
            - C1 (red), C2 (blue), C3 (gray) for a 3-color composite image lacking a green channel.
            - C1 (gray), C2 (cyan), C3 (magenta), C4 (yellow) for a 4-color composite image lacking a red, green, and blue channel.

    - Different "*Original File Format*" images may have different channel designations.
        - If the channel information of the image to be analyzed is unknown first open the merged composite multichannel 16-bit TIFF image or original file format image to be analyzed in FIJI/ImageJ.
            - Select: "*Image*" → "*Color*" → "*Channels Tool…*"
                - Toggle the various channels on and off to observed which color on the image corresponds to the respective channel.

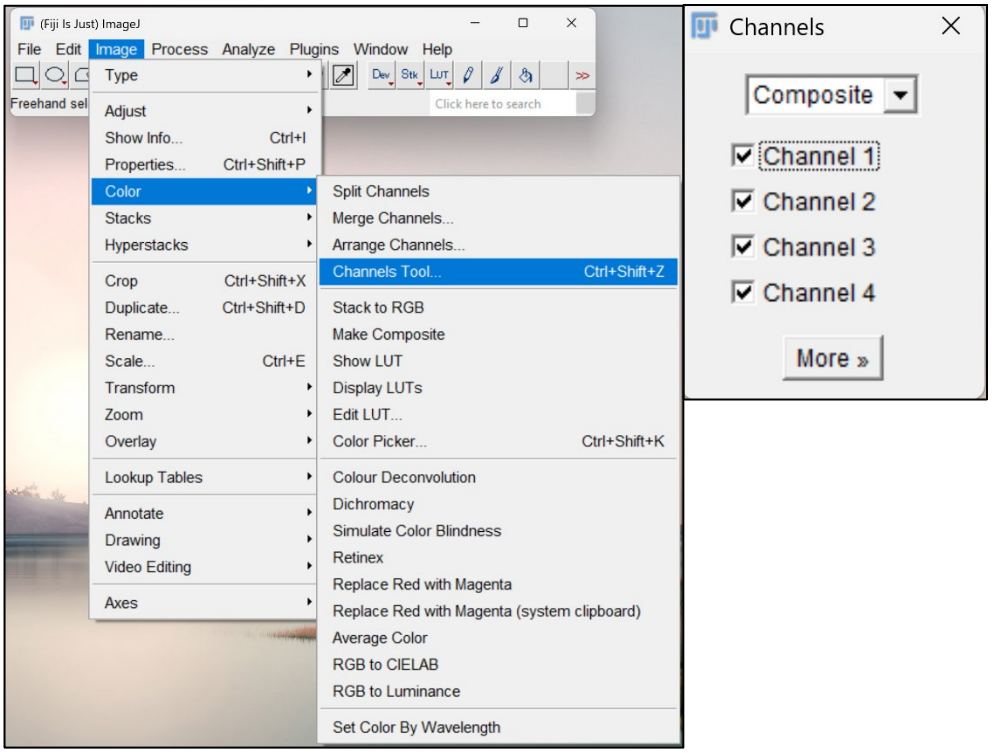


- MuscleBos assumes that the pixel-to-micron conversion factor is built into the metadata of the original file format or merged composite images to be analyzed or that images are manually calibrated prior to MuscleBos analysis.
  - If unsure whether the images to be analyzed are already calbibrated or rather require manual calibration open the image in FIJI and select: "*Analyze*" → "*Set Scale…*".
    - Confirm that the listed pixel-to-micron conversion matches that of both the specific microscope and objective used for imaging.
      - For example, a pixel-to-micron conversion of 2.822 pixels = 1 µm (0.354 µm per pixel) is the correct calibration factor for the Echo Revolution microscope which was used by us during MuscleBos creation and optimization when imaging with a 10 × objective
    - If the listed pixel-to-micron conversion reads "0" then the images are uncalibrated and manual entry of the appropriate calibration factor is required prior to MuscleBos analysis.
      - This is because the "*Original File Format*" Data format modes of the MuscleJ 1.0.2 source code and MuscleBos both set thresholds for object detection and report results in µm or µm$^2$ rather than pixels or pixels$^2$.

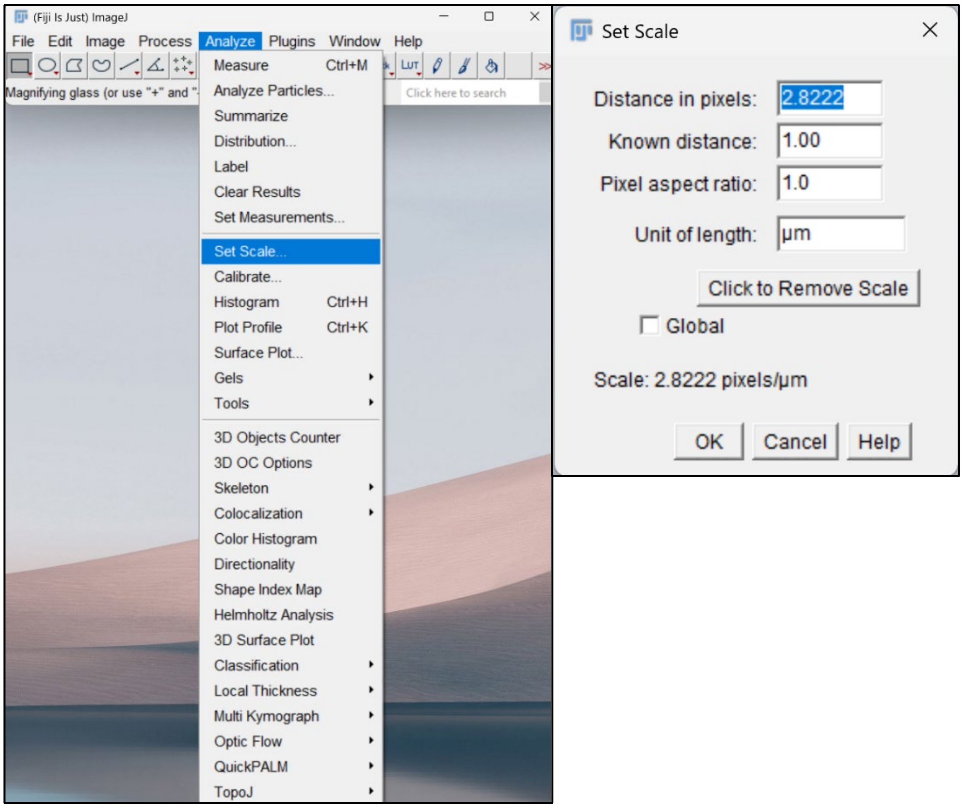


- Create a folder named “*Images*” as the source location of the image files to be analyzed and another named “*Results*” as a destination folder for the MuscleBos results output.
    - Place all the calibrated multi-channel composite TIFF (16-bit) or original file format images to be analyzed in the “*Images*” folder.
        - Ensure that no files other than the images to be analyzed are located within the “*Images*” folder.

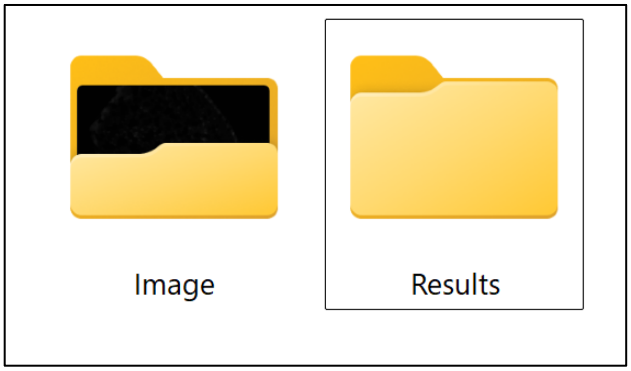


- Avoid using a single folder for both the source image input and results output.
    - Doing so introduced an error that limits the total number of images that can be analyzed in a single MuscleBos run.
        - In our hands this error is also observed in the source MuscleJ 1.0.2 code for an unknown reason.

**Step 2: Installation of the MuscleBos Plugin in the FIJI/ImageJ environment**

- Download FIJI from [https://imagej.net/Fiji/Downloads](https://imagej.net/Fiji/Downloads) and following the installation instructions. Alternatively, update the installed FIJI version to meet the minimum requirements.
    - MuscleBos was tested on FIJI version 2.14.0/1.52v with Java version: Java 1.8.0-66 (64 bits).
        - The minimum requirements are FIJI version 1.51e.

- Installation of MuscleBos
    - Download the “MuscleBos_V1-0.ijm” file onto your computer
    - Select “*Plugins*” → “*Install…*”
    - Locate the “MuscleBos_V1-0.ijm” file and save it in the FIJI “plugins” folder.
    - Restart FIJI to complete the MuscleBos installation.
        - The MuscleBos plugin should now appear in the FIJI “Plugins” menu.

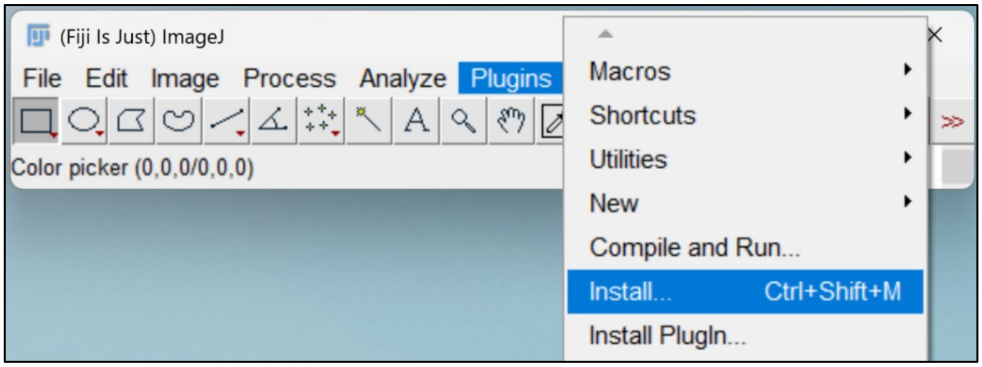


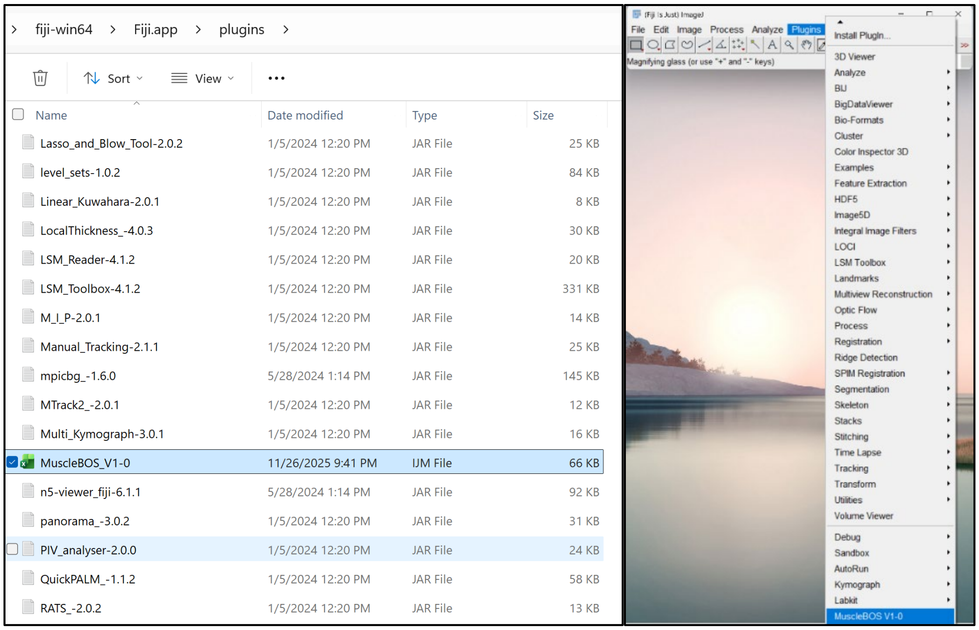

### Step 3: Running the MuscleBos plugin

- **Data Acquisition:**
  - **Volume selection**
    - **Single Z:** Chose this option for analysis of single Z-plane images.
    - **Z stack:** Choose this option if source file is a Z-stack image.
      - This will first create a maximum image projection of the source Z-stack images prior to analysis.

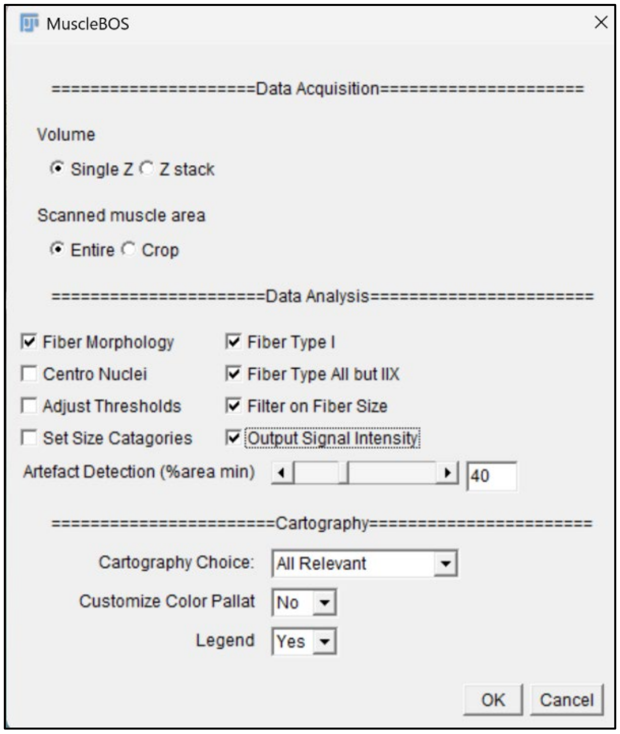


- **Scanned muscle area:**
    - **Entire:** Chose this option if the image to be analyzed is a stitched mosaic image encompassing the entire tissue cross-section.
      - This option is recommended if a compatible microscope equipped with a motorized stage and image stitching software is available.
    - **Crop:** Chose this option if the images to be analyzed are single field of view captured with a manual microscope or smaller cropped region of interest of the larger imaged entire tissue cross-section.
      - A series of additional filters are applied to track the maximum intensity of the fiber contours in the myofiber border stain channel.
        - This option has been retained from the MuscleJ 1.0.2 source code to enable use of MuscleBos for the analysis of manual capture of images of single fields of view or user defined cropped regions of a larger stitched mosaic image if desired.

- **Data Analysis:**
  - **Fiber Morphology:** Selecting this option utilizes a fluorescent stain of the myofiber boundaries (e.g., laminin, dystrophin, or wheat germ agglutinin) to detect and segment each individual muscle fiber detected within the image to be analyzed.
    - Selection of the "*Fiber Morphology*" option is mandatory to enable any additional more in depth "*Data Analysis*" options.

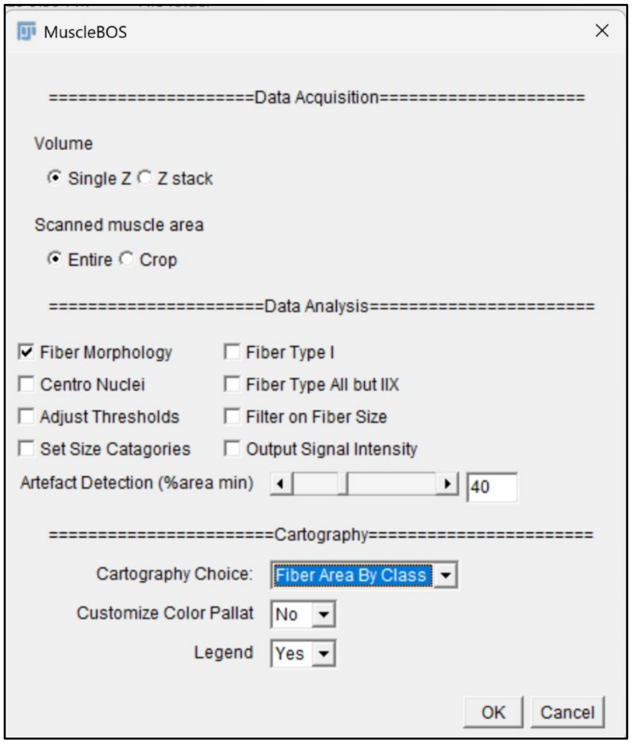


  - ***Channel Information:***
    - ***Fiber Shape*:** Select the appropriate channel of the image to be analyzed that corresponds to the muscle fiber border stain (e.g., laminin, dystrophic, or wheat germ agglutinin).

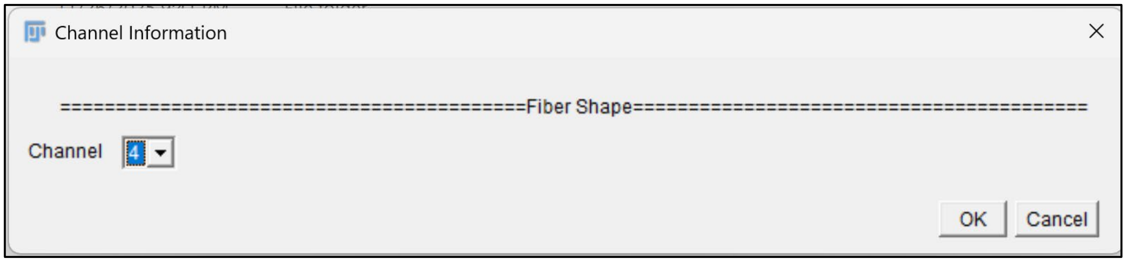


  - **Fiber Type I:** Selecting this option together with "*Fiber Morphology*" utilizes a fluorescent stain for slow-twitch type I myofibers (e.g., BA-D5) to classify the segmented muscle fibers as either "*Type I*" or not "*Type I*".
    - If "*Fiber Type I*" is selected without the "*Fiber All but Type IIX*" option selected, then the results outputs will characterize the segmented muscle fibers as either "*Type I*" fibers or ambiguous Type II fibers that will be designated as "*IIamb*".

- ***Channel Information:***
  - ***Fiber Shape:*** Select the appropriate channel of the merged composite image to be analyzed that corresponds to the muscle fiber border stain (e.g., laminin, dystrophic, or wheat germ agglutinin).

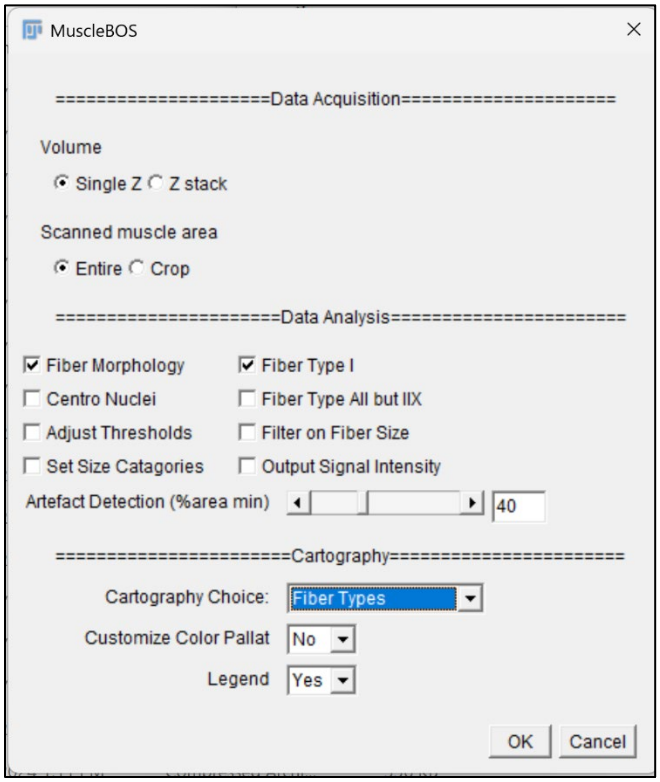


  - ***Type I:*** Select the appropriate channel of the merged composite image to be analyzed that corresponds to the fluorescent signal of type I muscle fibers (e.g., the BA-D5 primary antibody)

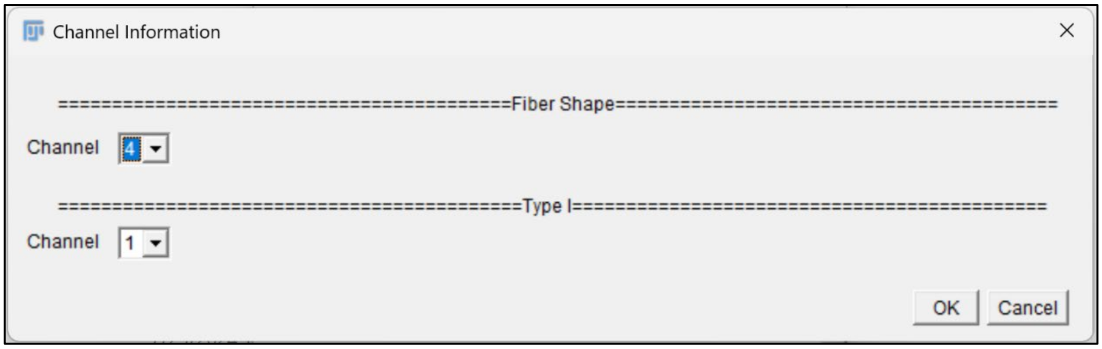


- **Fiber Type All but IIX:** Selecting this option utilizes a fluorescent stain of all myofibers except for type IIX fibers (e.g., BF-35) to classify fibers as either "*Type IIX*" (e.g., BF-35Neg) or "*Not Type IIX*" (e.g., BF-35Pos).
  - If the "*Fiber Type All but IIX*" option is chosen without also selecting "*Fiber Type I*" then segmented muscle fibers will be characterized as either "*Type IIX*" or "*Not type IIX*"

- ***Channel Information:***
  - ***Fiber Shape*: Select the appropriate channel of the image to be analyzed that corresponds to the muscle fiber border stain (e.g., laminin, dystrophic, or wheat germ agglutinin).
  - ***All but IIX:*** Select the appropriate channel of the merged composite image to be analyzed the corresponds to the fluorescent signal of all muscle fibers except for type IIX muscle fibers (e.g., the BF-35 primary antibody).

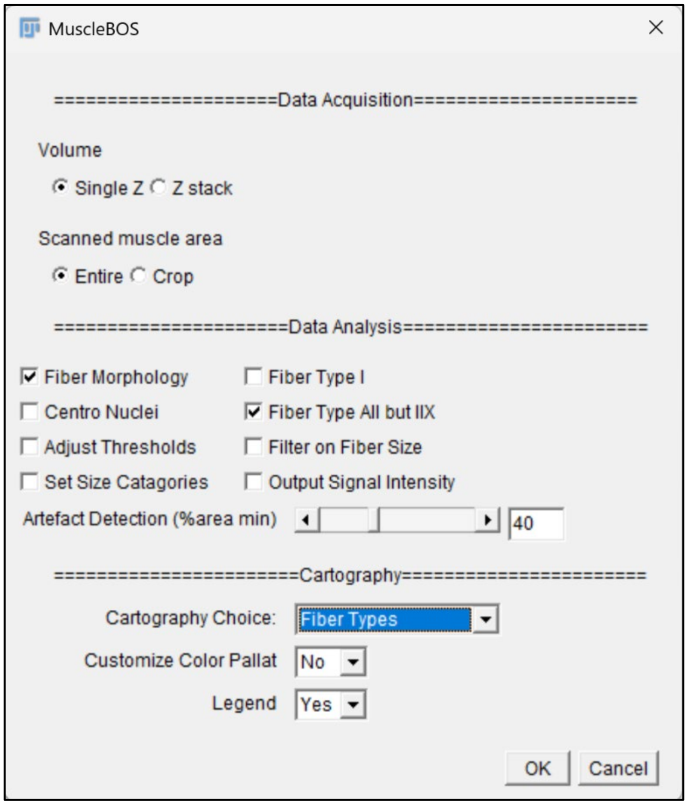


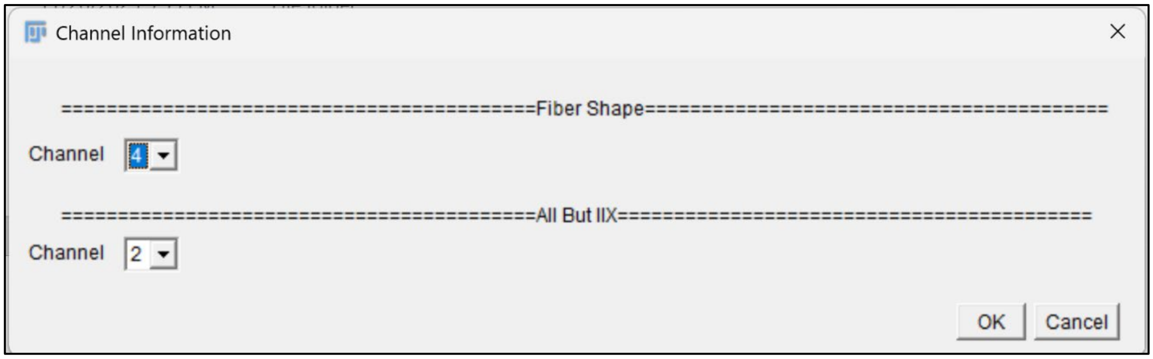


- **Combinations of Fiber Type identification:** Selecting the "*Fiber Type I*" option in combination with the "*Fiber Type All but IIX*" option utilizes fluorescent staining of type I fibers (e.g., BA-D5) in combination with all myofibers except for type IIX fibers (e.g., BF-35).
  - Segmented muscle fibers will be characterized as "*Type I*", "*Type IIA*", or "*Type IIX*".

- *Channel Information:*
  - ***Fiber Shape*: Select the appropriate channel of the image to be analyzed that corresponds to the muscle fiber border stain (e.g., laminin, dystrophic, or wheat germ agglutinin).

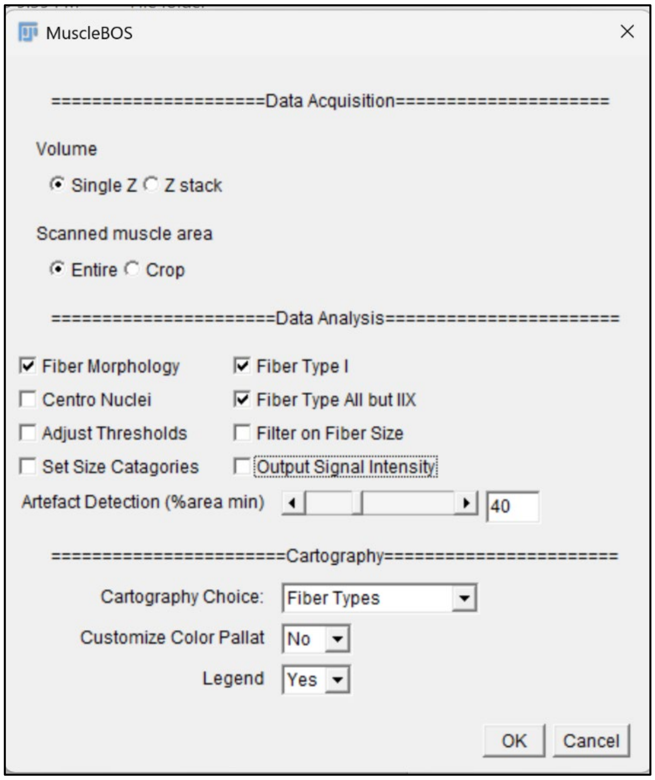


  - ***All but IIX:*** Select the appropriate channel of the merged composite image to be analyzed the corresponds to the fluorescent signal of all muscle fibers except for type IIX muscle fibers (e.g., the BF-35 primary antibody)

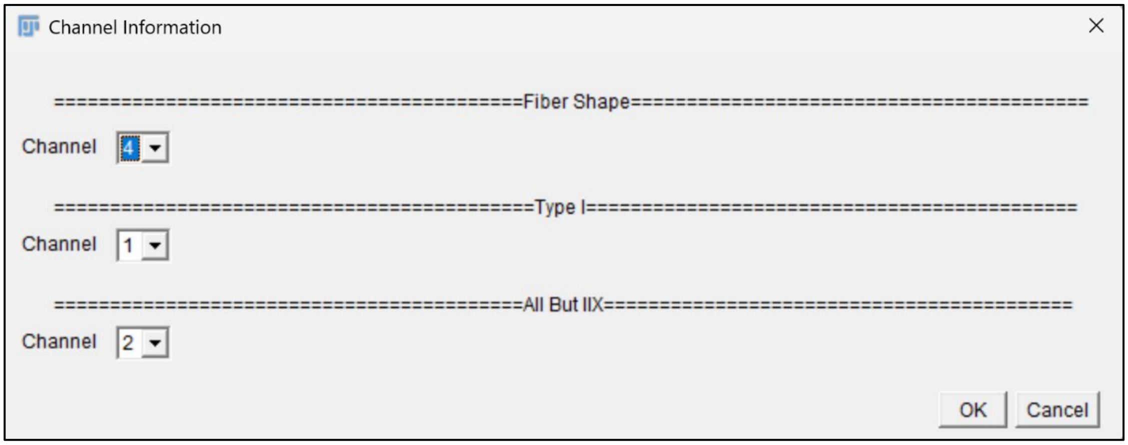


- **Centro Nuclei:** Selecting this option utilizing a fluorescent stain of the cell nuclei (e.g., DAPI) to count the number of peripherally and centrally located nuclei located within each segmented muscle fiber.
  - ***Note:*** In the current version of MuscleBos the "*Centro Nuclei*" option is incompatible with simultaneous selection of fiber type options including "*Fiber type I*" and/or "*Fiber type All but IIX*".
    - An error message will be displayed.

- ***Channel Information:***
  - ***Fiber Shape*****:** Select the appropriate channel of the image to be analyzed that corresponds to the muscle fiber border stain (e.g., laminin, dystrophic, or wheat germ agglutinin).
  - ***Nuclei*****:** Select the appropriate channel of the merged composite image to be analyzed that corresponds to the fluorescent signal of the cell nuclei (e.g., DAPI).

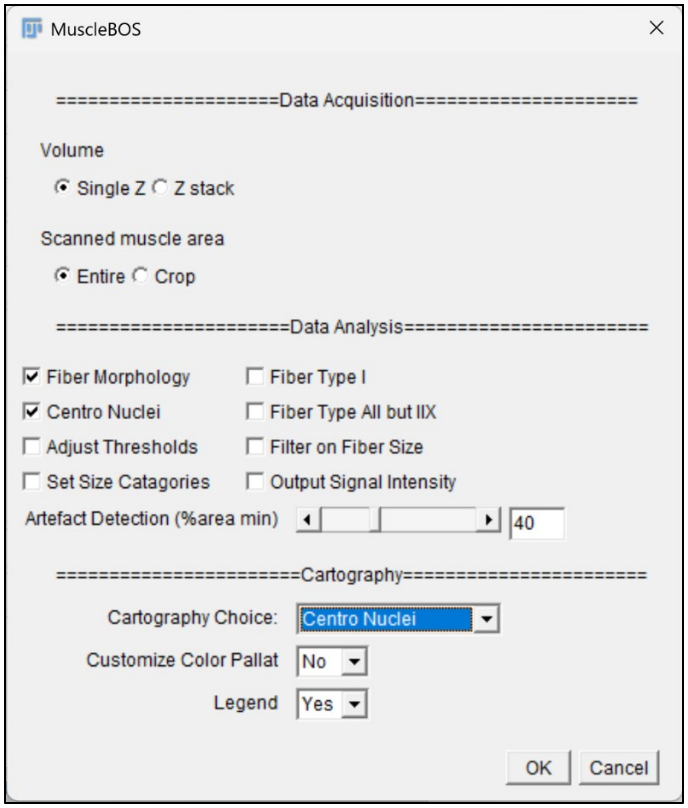


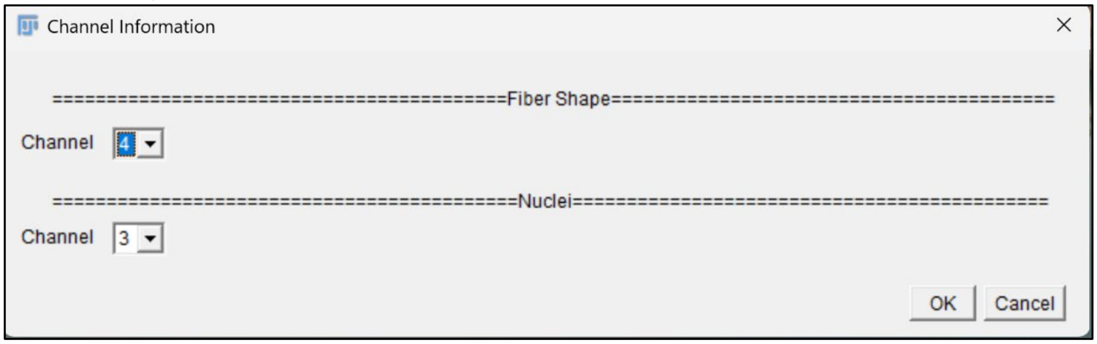


- **Adjust Thresholds:** Selecting this option enables additional options in the subsequent "*Channel Information*" dialog box that allow the user to set custom sensitivity thresholds for positive fiber type identification.
  - The adjust "*Fiber Type I*" threshold uses a single threshold defined as "X" standard deviations (SD) from the mean fiber type signal intensity to define fibers as either BA-D5Pos or BA-D5Neg
    - BA-D5Pos fibers are counted as "*Type I*".
    - BA-D5Neg fibers are counted as "*TypeIIAmb*"
      - If "*Fiber Type I* "is ran alone.
    - Or as either Type IIA or Type IIX
      - If "*Fiber Type I*" is selected together with "*Fiber Type All but IIX*".

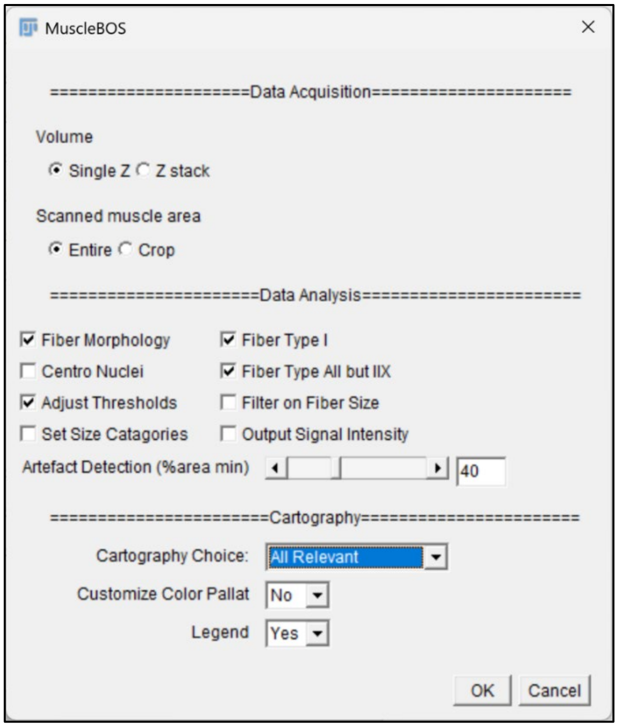


- The Adjust "*Fiber Type All but IIX*" option uses a both an upper and a lower SD threshold multiplier to characterize fibers as either BF-35Pos, BF-35Dim, or BF-35Neg
  - By default, the "*BF-35Pos*" and "*BF-35Dim*" fiber populations are pooled to obtain the "*Not type IIX*" population.
    - If running "*Fiber Type All but 2X*" option alone.
  - Or either Type IIA or Type IIX.
    - If "*Fiber Type I*" is selected together with "*Fiber Type All but IIX*").

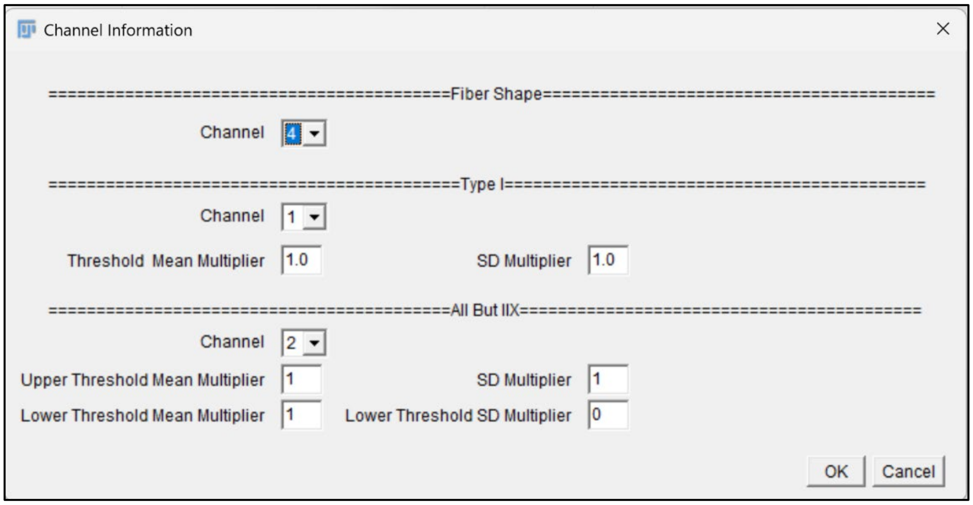

- **Set Size Categories:** Selecting the "*Set Size Categories*" option opens an additional dialog box allowing the user to set custom bins for fiber area frequency distribution analysis.
  - These bins are applied to both the "*Fiber Area*" cartography option and the frequency distribution data displayed in summary data output results sheet.

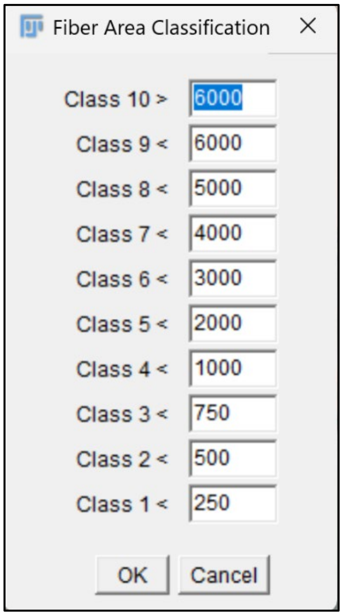


- **Filter on Fiber Size:** Selecting the "*Filter on Fiber Size*" option enables additional options in the subsequent "*Channel Information*" dialog box in which allow the user to set custom filters for minimal and maximal fiber CSA.
  - ***Note:*** By default, selecting the "*Filter on Fiber Size*" option will exclude unstained objects with a measured CSA <1000 µm from the "*Type IIAmb*" fiber population (if running the "*Fiber Type I*" option alone) or the "*Type IIX*" fiber population (if running both the "*Fiber Type 1*" and "*Fiber Type All but 2X*".

- **Output Signal Intensity:** Selecting the *"Output Signal Intensity"* option will save additional data columns in the detailed results output listing the signal intensity of each muscle fiber for each of the selected fiber type analyses.

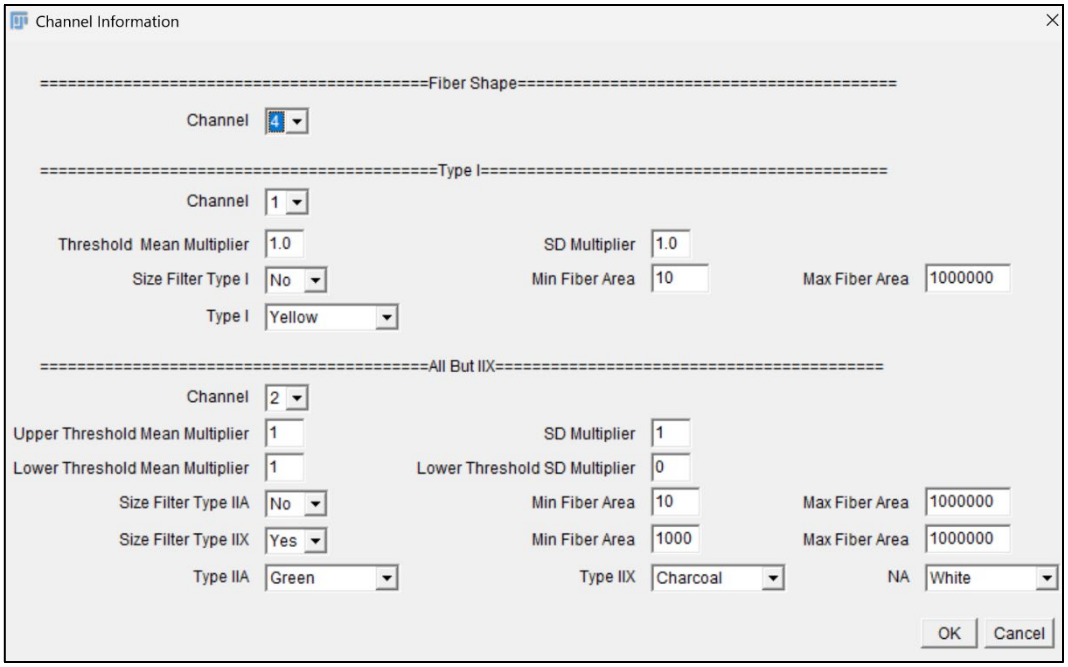

        - This allows users to use the raw data of fiber type staining signal intensities to characterize various fiber types if desired.

    - **Artefact Detection:** This option which is unchanged from the MuscleJ 1.0.2 source code allows the user to define an acceptable percentage of "artifact" (e.g., non-tissue area or poor staining).
        - If the quality of a given image falls below this threshold it is automatically flagged as an artifact and moved to the "Artefacts" results folder, thereby excluding this image from any further analysis.

- **Cartography:**

    - **Cartography Choice:** Selecting an option other than "*None*" will generate cartography files in the "*Results*" output folder.

        - **Fiber Area by Class:** Generates a cartography file of fiber ROIs color coded according to myofiber CSA.

        - **Fiber Types:** Generates a cartography file with muscle fiber ROIs color coded by muscle fiber type.
            - ***Note:*** In the current version of MuscleBos "*Fiber Type*" analysis is not compatible with "*Centro Nuclei*" analysis within a single run.

        - **Centro Nuclei:** Generates a cartography file with muscle fiber ROIs color coded according to the number of centrally located myonuclei.
            - ***Note:*** In the current version of MuscleBos "*Centro Nuclei*" analysis is not compatible with "*Fiber Type*" analysis within a single run.

        - **All Relevant:** Selecting this option will simultaneously generate multiple cartography files from a single MuscleBos run.
            - **Fiber Morphology + Fiber type:** Choosing "*All Relevant*" will output both "*Fiber Area by Class*" and "*Fiber Types*" cartography files.

    - **Fiber Morphology + Centro Nuclei:** Choosing "All Relevant" will output both "*Fiber Area by Class*" and "*Centro Nuclei"* cartography files.

- **Customize Color Palette:** Selecting "*Yes*" for this option will add additional options to the subsequent "*Channel Information*" dialog box that allow users user to choose custom color palettes for various "*Cartography*" results file outputs.
